# Building a National HIV Cohort from Routine Laboratory Data: Probabilistic Record-Linkage with Graphs

**DOI:** 10.1101/450304

**Authors:** Jacob Bor, William MacLeod, Katia Oleinik, James Potter, Alana T. Brennan, Sue Candy, Mhairi Maskew, Matthew P. Fox, Ian Sanne, Wendy S. Stevens, Sergio Carmona

## Abstract

**Background:** Chronic disease management requires the ability to link patient records across multiple interactions with the health sector. South Africa’s National Health Laboratory Service (NHLS) conducts all routine laboratory monitoring for the country’s national public sector HIV program. However, the absence of a validated patient identifier has limited the potential of the NHLS database for epidemiological research, policy evaluation, and longitudinal patient care. We developed and validated a record linkage algorithm, creating a unique patient identifier and enabling analysis of the NHLS database as a national HIV cohort. To our knowledge, this is the first national HIV cohort in any low-or middle-income country.

**Methods.:** We linked data on all CD4 counts, HIV viral loads (VL), and ART workup laboratory tests from 2004-2016. Each NHLS laboratory test result is associated with a name, sex, date of birth (DOB), gender, and facility. However, due to typographical and other errors and patient mobility between facilities, different patient specimens may be associated with different sets of identifying information. We developed a graph-based probabilistic record linkage algorithm and used it to construct a unique identifier for all patients with laboratory results in the national HIV program. We used standard probabilistic linkage methods with Jaro-Winkler string comparisons and weights informed by response frequency. We also used graph concepts to guide the linkage in determining whether a cluster of patient specimens could plausibly reflect a single patient. This approach allows matching thresholds to vary with the density of the network and limits over-matching.

To train and validate our approach, we constructed a quasi-gold standard based on manual review of 59,000 candidate matches associated with 1000 randomly sampled specimens. These data were divided into training and validation sets. Domain weights and graph parameters were optimized using the manually matched training data.

To evaluate performance, we calculated the probability that a true match was correctly identified by our algorithm (sensitivity, Sen) and the probability that a match identified by our algorithm was truly a match (positive predictive value, PPV) in the manually-matched data. We also assessed validity in the full cohort using proxies for under-and over-matching and assessed sensitivity vis-à-vis national identification numbers and patient folder numbers, which were available for a sub-set of records. We compared the performance of our algorithm for exact matching and a prior identifier that had been developed by the NHLS Corporate Data Warehouse.

**Results.:** As of December 2016, the NHLS database contained 117 million patient specimens with a CD4, VL, or other laboratory test used in HIV care. These specimens had 63 million unique combinations of patient identifying information. From these data, our matching algorithm identified 11.6 million unique HIV patients who had at least one CD4 count or VL result. These patients 70.9 million total specimens, with a median of 3 specimens per patient (IQR 1 to 8). Sensitivity and PPV of the algorithm were estimated to be 93.7% and 98.6% in manually-matched data, compared to 64.1% and 100.0% for the existing NHLS identifier. We estimated that in 2016 there were 3.35 million patients on ART and virologically monitored, similar to the National Department of Health estimate of 3.50 million.

**Conclusion.:** We constructed a South African National HIV Cohort by applying novel graph-based probabilistic record linkage techniques to routinely collected laboratory data, with high sensitivity and positive predictive value. Information on graph structure can guide record linkage in large populations when identifying data are limited.

## 1. INTRODUCTION

### 1.1 The public health rationale for record linkage

Management of chronic diseases like HIV requires the ability to link patient records across multiple interactions with the health sector. Record linkage presents a challenge in large populations where unique patient identifiers (e.g. Social Security Numbers) are not systematically recorded and where other identifying information is limited. This scenario is common in many developing countries faced with a growing burden of chronic disease, yet where health systems were principally designed to provide acute and preventive care.^1^ We develop a scalable, graph-based approach to probabilistic record linkage, and apply it to the complete laboratory records of South Africa’s national HIV treatment program, the largest in the world.

Interest in record linkage methods among health researchers has increased with the proliferation of “big data”, i.e. data generated through routine interactions rather than for research purposes, and with better access to the computational resources needed to link these data. When no unique identifier (e.g. social security number, national ID number, passport number) exists, other identifying data can be used to assign records to individuals probabilistically.^2,3^ Accurate record linkage is a key step in transforming “big data”, including clinical and administrative data, into databases usable for epidemiologic research, program evaluation, and longitudinal monitoring of patients with chronic conditions.

### 1.2 Record linkage within a single database

This paper focuses on the linkage of records within a single database, also known as deduplication, disambiguation, or entity resolution.^2^ There are several key features of our application, which distinguish it from other record linkage problems.

1. <u>The task is to identify all records associated with the same patient.</u> The data consist of unlinked patient specimens. Each specimen is associated with some identifying information, which may be reported with error. Multiple laboratory tests may be conducted on each specimen, and these tests will have identical identifying information.
2. <u>There is no gold standard listing of patients</u>. The deduplication process will recover a set of patients in the data, but there is no master list of true patients against which to validate. Furthermore, the true underlying number of patients giving rise to the data is not precisely known. (There are however external estimates against which to compare as a validity check.)
3. <u>A patient may give rise to any number of specimens with similar but non-identical identifying information</u>. As a result, a patient specimen could be correctly linked to any number of other specimens. That number is unknown, although though there is a plausible range.
4. <u>A single record cannot belong to multiple patients.</u> This implies that transitivity holds. If A links to B and B links to C, then A and C are implicitly linked and attributed to the same patient, in contrast to 1:1 linkage across databases.
5. <u>All valid results should be assigned to patients</u>. After removing invalid specimens, i.e. those associated with research studies or quality assurance, all other specimens arise from real patients. Because the goal is to match all specimens to patients, we do not throw away records that could match to multiple patients, unless identifying information in key domains is missing or the result was invalid.
6. <u>The database is large</u>. Our linkage focuses on 116 million CD4 count, viral load, and other plausibly-HIV-related laboratory tests collected in South Africa’s public-sector HIV program since 2004, corresponding to 63 million unique sets of identifying information.

Probabilistic record linkage dates to the mid-20^th^ century. Newcombe (1959) showed that record linkage can be framed as an optimization problem, where the goal is to minimize both over-matching and under-matching errors.^5^ Over-matching occurs when results that were not generated for a patient are attributed to that patient, leading patients to be falsely combined. Under-matching occurs when results that were generated by a patient are not attributed to that patient, leading to the appearance of additional “patients” in the dataset. A more liberal matching rule will reduce under-matching, leading to greater Sensitivity, but will result in more over-matching and lower Positive Predictive Value (PPV). (Conversely, a more conservative matching rule will increase PPV, but reduce Sensitivity.) Newcombe proposed what has become the traditional approach to this problem: compare records on a range of characteristics; generate a similarity score for each characteristic and combine into a total similarity score; and then choose thresholds denoting whether the link is “considered a match”, “not considered a match” or “held for manual review.” Fellegi & Sunter (1969) derived a formula for the optimal similarity score, which incorporates empirical information on response frequencies within domains and on random (e.g. typographical) error rates.^6^ For example, a match on a rare name is less likely to occur by chance than a match on a common name and thus receives greater credit in the similarity score.

Record linkage becomes more difficult in large datasets. First, it may be impossible to compare all observations with all other observations. Strategies such as blocking (restricting comparisons to observations that match on some characteristics) are needed to reduce the number of comparisons to be scored, resulting in a trade-off between computational efficiency and the possibility that some true links are not considered. Second, once potential matches are scored, the traditional approach of using manual review to resolve uncertain matches may be impossible due to the large number of candidate matches to evaluate. Third, the probability of over-matching increases with the size of the underlying population, and hence the costs of foregoing manual review increase. The larger the size of the population, the greater the chances that a random (e.g. typographical) error will falsely link specimens from two different people, A—B. When such a false dyad emerges, its chances of acquiring an additional false match are approximately twice what they would have been in the absence of the initial linkage error. Without additional identifying information, over-matching in big data sets can lead false links to amplify into very large clusters, ultimately leading to the false linkage of individuals with completely different sets of identifying information through several degrees of separation.

### 1.3 Using graphs to guide record linkage

We propose a graph-based solution to address the scalability problem.^2,7–12^ We define nodes as unique sets of patient identifying information as recorded in the NHLS database. We define weighted edges as the scored comparisons between these nodes, with scores calculated using a modified version of the Fellegi-Sunter approach.^6^ The complete laboratory database can be interpreted as a very large graph (i.e., network) defined by these nodes and edges. Individual patients are represented by connected components (or clusters) of the graph. Previous approaches have used clustering algorithms in the process of entity resolution,^11,13^ or have used graph concepts to screen for possible linkage errors for manual review^7,8^. However, most existing off-the-shelf record linkage packages do not use graph-concepts to guide the linkage.

We use information on the graph-structure of individual clusters to help determine whether the cluster represents a single patient. Our approach is based on a simple heuristic: a cluster cannot represent a single patient if the two most dissimilar nodes in the cluster do not reflect a single patient. In effect, our approach replaces the threshold rule used to determine the existence of edges in the traditional Fellegi-Sunter approach with a threshold rule on the weighted diameter of the cluster, i.e. the shortest weighted path between the farthest points. Unweighted diameter was shown to outperform other graph metrics (number of bridges, density of graph) in identifying false clusters in Australian administrative health records.^7^ Weighted diameter incorporates even more information on the strength of the ties and better captures the likelihood that the cluster represents a single patient.

This graph-based approach to record linkage exploits information contained in the graph structure not typically used in traditional linkage approaches. First, it brings in useful information on the size and shape of the cluster. The data generating process for errors in identifying information is complex, but follows some known rules, which can be incorporated into the scoring function and yields predictable shapes and sizes. For example, long chains of observations in which A links only to B, B links only to C, and C links only to D, etc., are unlikely. On the other hand, it is not uncommon to have “missing edges” in a cluster; e.g. B may contain information (an English and Zulu name) that provides a “key” correctly linking A and C even though the name dissimilarity between A and C implied no direct link.

Second, the graph-based approach incorporates useful information about the “neighborhood” of the cluster, i.e. other records that may be similar but come from distinct patients. Clusters in sparser areas of the graph are less likely to falsely combine with data from other patients and therefore we can allow for lower scored edges and greater linkage sensitivity. By contrast, higher standards are needed in denser areas of the graph to prevent false linkage, e.g. of names such as James, Jones, Jan, Jason.

By incorporating these sources of additional information, the graph-based approach has several practical benefits for linkage of large datasets:

a. The graph-based approach reduces the need for manual review of borderline cases.
b. The graph-based approach limits the impact of overmatching in large datasets, flagging and breaking up implausible clusters before they become very large.
c. Weighted diameter provides an intuitive justification for why clusters should be considered patients and why others should not, based on the similarity of the most dissimilar records.
d. The threshold for the weighted diameter can be adjusted as an algorithm input parameter. In doing so, the analyst can identify unstable clusters that are sensitive to the threshold choice (typically those in denser regions of the graph) and stable clusters that are less sensitive to the threshold choice (typically those in sparser regions). At the data analysis stage, the subset of stable clusters can be used to assess the robustness of estimates in patients where linkage errors are minimized. Unstable clusters can also be reviewed manually to verify choice of threshold.

We developed this graph-based probabilistic record linkage algorithm in the context of a collaboration with South Africa’s National Health Laboratory Service (NHLS) to support NHLS in monitoring and evaluating the country’s national HIV program. We therefore illustrate the performance of the approach with this real-world application. NHLS conducts all laboratory monitoring for South Africa’s national HIV program. By creating a validated unique identifier in the NHLS database, we construct what is – to our knowledge – the first national HIV cohort in any low-or middle-income country.

The paper proceeds as follows. Section 2 describes the NHLS database and describes the development of a manually matched training and validation set. Section 3 presents the record linkage algorithm, with sub-sections on pre-processing, search, scoring, graph-based entity resolution, and computational performance. Section 4 presents validation results and some summary statistics on the resulting dataset. Section 5 concludes.

## 2. DATA

### 2.1. South Africa’s National Health Laboratory Service Database

South Africa has the world’s largest HIV burden at 7.1 million people infected, representing a fifth of the HIV-infected population worldwide.^14^ The country also has the world’s largest HIV treatment program with about 4.4 million people on antiretroviral therapy (ART) in 2018 It has been estimated that 91% of patients on HIV treatment are receiving ART in the public sector.^15^ With ART, people living with HIV (PLHV) can have long, healthy, and productive lives.^16–18^ ART also reduces the chances of onward transmission of the virus.^19,20^ As a result of South Africa’s large investment in HIV treatment, population life expectancy has increased by over a decade in some regions.^21^ However many PLHV are not yet on therapy, and the country has introduced new policies to significantly expand treatment coverage^22^ with the goal of reducing transmission^23^ and ending the epidemic.

HIV disease progression and treatment success are monitored primarily through regular laboratory tests: CD4 counts to assess immune function and viral loads (VL) to assess the concentration of the virus in the blood. NHLS is the sole provider of diagnostic and monitoring pathology services for those accessing HIV care in the public sector and has done so since program inception in 2004 (with the exception of one province – KwaZulu-Natal – which joined the NHLS in 2010.) Although guidelines have changed periodically since 2004, a CD4 count has always been conducted following HIV diagnosis and either CD4 counts or VLs have been conducted at least annually to monitor treatment efficacy. As of December 2016, the NHLS Corporate Data Warehouse (CDW) contained records of 32.5 million CD4 counts and 20.1 million VL since 2004 conducted on 46 million patient specimens. In addition to CD4 counts and VLs, NHLS provides clinics with laboratory support for other laboratory tests used in HIV monitoring and treatment decisions – Alanine Aminotransferase (ALT), Hemoglobin, Cryptococcal Antigen, Creatinine Clearance, and HIV PCR/Elisa results – and our data included 102 million of these tests on 71 million specimens. (These tests are also used for patients without HIV.) In total, the NHLS database included over 117 million patient specimens with over 154 million tests conducted that could possibly be related to HIV care between 2004 and 2016.

The CDW contains three sources of data: patient demographics, laboratory test results, and facility characteristics. The laboratory results data in the NHLS database are comprehensive and accurate. Specimens are collected at public sector clinics and hospitals, and are analyzed either at that facility or at one of several NHLS reference laboratories. Data on patient demographics, facility, and laboratory results are captured into an electronic laboratory information system (LIS). Each specimen is assigned a unique “Episode Number” (Episode_No), which is the link between the patient demographics, i.e. name, gender, and date of birth, location of the referring facility, and the results of any test requested. Results are delivered to facilities for patient care via: paper hard copy, SMS printing, an online query system, and/or telephonically. Data in LIS are then checked and are transferred to the NHLS CDW database in near-real time. Because the data are obtained directly from the LIS, they are less vulnerable to gaps in clinical record keeping at the facilities. As a result, the NHLS database provides a more complete representation of laboratory testing than South Africa’s electronic health monitoring database (TIER.Net), as many facilities only report laboratory test results to TIER.Net if they have been copied manually into patient charts and later extracted from the patient charts into TIER.Net.^24^

### 2.2 Need for a Validated Unique Identifier in the NHLS Database

A key limiting factor in the NHLS Database is the variety and accuracy of the identifying information collected. The demographic information fields available on the laboratory requisition form include national ID number, patient folder number, surname, first name, sex, date of birth (or age), physical address and patient telephone number. This information is collected from the patient and then captured on the LIS at the NHLS registration site. Many of the demographic fields can be incomplete and collection and transcription errors are common.

Therefore, the major limitation of the NHLS data is the lack of a validated unique patient identifier to enable linkage across all laboratory test results associated with a single patient. The NHLS data are curated at the level of the test result. From 2004-2016, national ID numbers were collected for only 2% of specimens. And despite the high quality of the data, there remains substantial variation in the patient identifying information associated with patient specimens. For example, our extract of 117 million specimens contained 62.8 million unique sets of identifying information. Yet the population of South Africa is only 56.6 million.^4^

Variability in names arises from several sources. Predictable sources of variability include: typographical errors (Alex vs. Alwx), hearing errors (Alex vs. Alice), nicknames (Alex vs. Alexander), translations (Mpho vs. Gift), first/last name inversions, use of multiple first and middle names, and abbreviations (VD vs. van der). Other variation arises from extraneous information, e.g. titles, prefixes, redundant initials, non-alphabetical characters, which can be addressed in pre-processing the data. Still other variations are less predictable: e.g. English vs. local language names (Beatrice v Nonhlanhla), name changes at marriage, and other name changes. Finally, sometimes a name is simply unknown (e.g. No Name, Mother of), or may not exist, in the case of some neonates (e.g., Baby, Twin).

Variation in dates of birth can arise from typographical and hearing errors, from month-day inversions, from false reports or misremembering, or through provision of an age rather than an exact date of birth. Gender may be listed incorrectly due to typographical error, due to the large proportion of androgynous names in this setting and may also change if some patients transition to other genders. Finally, for all of these domains, patients may deliberately obfuscate their identifying information. For example, patients may provide false information to avoid discovery in an environment of HIV stigma, to access care if they are not citizens, or to “shop” for care at other facilities.

We set out to create a validated unique patient identifier in the database of all HIV-related test results, using existing demographic information recorded for each specimen: first name, last name, date of birth, gender, province, and facility. The development of a valid unique patient identifier would have several important implications. First, a unique patient identifier would enable de-duplication, in order to achieve accurate reporting of aggregate trends, such as the number of patients in pre-ART care and the number of patients on ART and monitored for VL. Second, it would enable monitoring of longitudinal concepts such as CD4 recovery, virological failure, and retention in care, enabling identification of low-performing “hot-spots” and high-performing model facilities. Third, a unique patient identifier would enable the construction of a National HIV Cohort, which can be used for longitudinal epidemiological analysis and evaluation of policies and programs. Finally, if sufficiently high accuracy were attained, a unique patient identifier could be integrated into electronic medical records systems, offering providers at any networked facility access to a patient’s complete history of laboratory test results and improving chronic disease management in mobile patient populations.

We note that NHLS’s CDW previously developed a linkage algorithm, however its validity is unknown. Early analysis of the CDW unique identifier suggested evidence of overmatching as there were some implausibly large clusters. There was also evidence of under-matching as the algorithm identified 18.5M unique HIV patients, an implausibly large number of patients given that just over 7M South Africans are currently HIV-infected and about 2.5M have died since 2004.^25,26^ We evaluate the performance of the graph-based linkage algorithm we developed alongside the CDW unique identifier as well as a simple “exact match” on name, gender, and date of birth.

### 2.3. Developing a manually matched quasi-gold standard

#### Original manual-matching exercise

Record linkage can be substantially improved with the existence of training data to optimize the algorithm. Additionally, record linkage exercises ideally validate the results against a gold standard. In the case of the NHLS database, no gold standard dataset exists that captures the potential flow of patients across different sites within South Africa’s public-sector health system.

To train and evaluate the algorithm, we constructed a manually-coded quasi-gold standard dataset. We randomly selected 1000 patient specimens from the full database of 30.4M specimens with an CD4/VL result available in Fall 2014. For each of these 1000 “index” specimens, we started with a very liberal (high sensitivity) early version of our matching algorithm and generated candidate matches from the full 30.4M specimen database. We identified an average of 59 candidate matches per index specimen (range 0, 838). Four trained research assistants (RAs) manually evaluated these 59,000 candidate matches for match quality on a 4-pt scale: 1 = almost certainly not a match, 2 = plausible match, 3 = probable match, 4 = almost certain match. After an initial training period to harmonize evaluations, each candidate match was graded twice by separate RAs. After all matches were graded, we held a refresher training session. Then a third RA reviewed all candidate matches for which there was disagreement between the first two RAs and determined a final match quality. RAs had access to additional – though highly incomplete - information on patient addresses, test dates, and national ID numbers, which could sometimes be used to improve the manual match. Finally, to limit over-matching, we conducted a targeted re-review of all “patients” that moved between provinces multiple times, were reported as both male and female, or had common names. We considered all 3s and 4s as “matches” and all 1s and 2s as “not matches”. The result of this exercise was a manually matched quasi-gold standard dataset consisting of all laboratory results linked to the same patient as a random sample of 1000 laboratory results.

We refer to the manually-coded data as a “quasi-” gold standard because they reflect our best human assessment. The RAs often had to make judgments amidst uncertainty as to whether particular candidates were matches or not. RA intuition was “tuned” through training and team discussions of difficult cases. To assess whether the RAs were identifying approximately the right number of matches, we conducted a back-of-the-envelope calculation: We used the distribution of numbers of specimens per patient identified by the RAs as “matches” to estimate the number of true patients that gave rise to the total number of specimens in the NHLS database with CD4/VL tests. Our estimate of 10.2M people with at least one CD4/VL test (in the realm of plausibility) suggested that the RAs were linking a reasonable number of results to patients and were neither too strict nor too lax in determining matches.

#### Training and validation sets

The same dataset cannot be used both to guide choices about the algorithm and then to evaluate the performance of the algorithm, since resulting estimates will be biased. We therefore randomly divided the gold standard dataset of 1000 specimens - and 59,000 manually-scored comparisons - into two sub-sets which can be considered independent samples: 489 specimens (and their scored candidate matches) were used as training data; and 495 specimens (and their scored candidate matches) were reserved as a validation set, to be set aside and used only to evaluate the performance of the final, optimized algorithm. (Eleven of the index specimens in the training set and 5 of the index specimens in the validation set were found to be invalid records after sampling.) The training and validation sets are summarized in **Table 1**.

**Table 1.** Description of manually coded “quasi-gold standard” dataset

|  | **Index**<br>**specimens, n** | **Manually-coded**<br>**“true matches”,**<br>**original; revised** | **Manually-coded**<br>**“true non-matches”,**<br>**original; revised** |
| --- | --- | --- | --- |
| Training set | 489 | 3678; 3899 | 20,453; 20,232 |
| Validation set | 495 | 3840; 4284 | 19,858; 19,432 |

Note: True matches and true non-matches were determined based on manual review. The revised manually coded data include some candidate matches not identified in the original review, so the total numbers of candidate matches differ. Methods for developing the original and revised manually-coded data are described in the text. We note that different RA’s coded the training and validation sets which were carried out as separate exercises. A larger number of matches were identified for the validation set relative to the training set, which likely reflects that the second RA team was slightly more liberal in determining what counted as a match.

#### Reducing noise in the manually-matched data

There were a number of close cases in which the RAs reported that they simply had to make their best guess. Additionally, despite an extensive training period and at least two evaluations of each comparison, there were differences across RAs in the distributions of scores and there may have also been within-RA variation in coding as a result of temporal differences in alertness or attention to specific patterns in the data. These factors may have led to random errors in the manually-coded data that were not present in the underlying ground truth data.

When a gold standard is measured with error, this leads to sub-optimal training leading to a reduction in performance. It also leads to a reduction in perceived performance in relation to the validation set because the random component is by definition unpredictable. As we refined our algorithm and compared this improved algorithm to the training data, we found that many of the discrepancies between the manually-and algorithm-coded results were most likely caused by manual errors in the RA-coded data. Additionally, the improved algorithm identified some new candidate edges that had not been scored in the original manual review because the refined algorithm had improved sensitivity relative to the version initially used to identify candidate matches.

We therefore undertook a process to update the training and validation sets, following published methods for test validation when the gold standard is measured with error.^27^ We implemented the improved algorithm, setting the tuning parameters to achieve high sensitivity. This allowed us to identify new candidate matches that were not identified by the original algorithm for further review. Two RAs then re-assessed the following: all (validation) or a random subset (training) of candidate matches in which both the coders and the algorithm agreed it was a match; all candidate matches in which there was disagreement; all candidate matches not scored by the original coders; and a random subset of candidate matches in which the algorithm and original coders agreed there was no match. Because this last category constituted the vast majority of the manually-matched dataset, the task for the RAs was substantially reduced in relation to the original review. The RAs were blinded as to the original computer and coder assessments. A member of the research team, also blinded, then re-evaluated all cases of discordance between the two RAs and between the RAs and the original coders, and conducted additional spot checks. The updated manually-coded data were used for all subsequent training of the algorithm. (We note that the re-review of the training and validation data were conducted as separate exercises by separate RA teams, which may have led to some differences between the revised training and validation sets.)

### 2.4 Evaluating Sensitivity and Positive Predictive value in quasi-gold standard data

In training and validating the algorithm with respect to the manually-matched quasi-gold standard, we focused on two parameters commonly used in evaluating record linkage^2^:

1. Sensitivity, i.e., the proportion of manually-coded true matches that were correctly identified by the algorithm, and
2. Positive Predictive Value (PPV), i.e., the proportion of matches identified by the algorithm that were manually-coded as true matches.

Sensitivity and PPV are also known as recall and precision, respectively, in the computer science literature. These parameters are defined at the candidate match level (not at the patient level). They are calculated as follows: randomly choose a patient specimen; then identify all other specimens associated with the same patient according to the algorithm vs. according to the quasi-gold standard. Because the number of non-matches is so large, Specificity and Negative Predictive Value will nearly always be close to 100% and therefore are not useful for training or validation.

All training was conducted with the best available manually-matched data. Initial training was conducted using the original version. Later training was conducted using the revised manually-matched data, which was believed to be closer to ground truth.

In our validation exercise, we report Sensitivity and PPV both with respect to the original manually-coded test data as well as compared to the revised manually-coded test data. Both versions are reported for transparency. The revised manually-coded data were heavily scrutinized through additional rounds of review and are believed to be closer to truth. Ignoring these improvements, our Sensitivity and PPV estimates with respect to the original data are likely to be biased downwards. On the other hand, using the algorithm to guide the manual matching has potential to lead to biased evaluation of algorithm performance vis-à-vis those data. In order to minimize potential for bias in the second round of manual review, we blinded the reviewers as to how the algorithm coded a particular candidate match. Additionally, to obtain accurate estimates of Sensitivity and PPV from the revised validation set, it is necessary to assess and adjust for the possibility that there were cases of false agreement between the computer and manual coders, not only cases of false disagreement. We report revised estimates of Sensitivity and PPV using published formulas, which account for this possibility.^27^

## 3. A GRAPH-BASED RECORD LINKAGE ALGORITHM

### 3.1 Overview of the approach

Record linkage has become an increasingly common activity for governments and private sector organizations as the extent of administrative and other big data has increased and as computing power to conduct record linkage has improved. When there is no unique identifier, probabilistic or “fuzzy” matching techniques can be employed to develop an identifier. The central task in fuzzy record linkage is to create a unique identifier that simultaneously minimizes both over-matching (falsely combining records that should remain separate) and under-matching (falsely separating records that should combined). Our chosen approach was based on a review of existing best practices in the literature and consultation with several authorities on record linkage.

Our approach was guided by several principles that we committed to at the outset:

1. We should attempt to capture different sources of systematic errors common in South Africa, including: specific types of data entry errors, use of nicknames and multiple names, and uncertainty about dates of birth;
2. We should use ONLY demographic information and should avoid using clinical information (e.g. CD4 values) to match, as including such information would bias our results towards the patterns in health outcomes that we seek to measure.
3. We should use fuzzy matching methods applicable to a setting with 11 national languages. This ruled out “Soundex” type methods, which exploit similarities in how words or syllables sound and are thus language specific.
4. We should exploit the fact that some names and dates of birth are more common than others and may be more similar to other names than others.
5. Our methods should be scalable to the NHLS’s very large datasets. Record linkage is more difficult the larger the number of records, since there is much greater potential for over-matching. Additionally, larger datasets require more computing resources and a blocking strategy which limits comparisons.
6. Any resulting algorithm must be validated and the extent of over-matching and under-matching error reported in an unbiased way.

Our goal was not perfection, but the “best” unique identifier that we could come up with and clear, unbiased reporting on the quality of the identifier.

Our record linkage method consisted of four steps (**Figure 1**): pre-processing, search for edges, scoring edges, and graph-guided entity resolution. The methods were implemented using Boston University’s Shared Computing Cluster (SCC), which hosts secure data and meets standards for dbGaP compliance, includes many statistical programmes, and consists of a network of high speed, multiprocessor computers. We received key technical support from Senior Data Scientist, Katia Oleinik, who supports researchers using the SCC.

**Figure 1.**
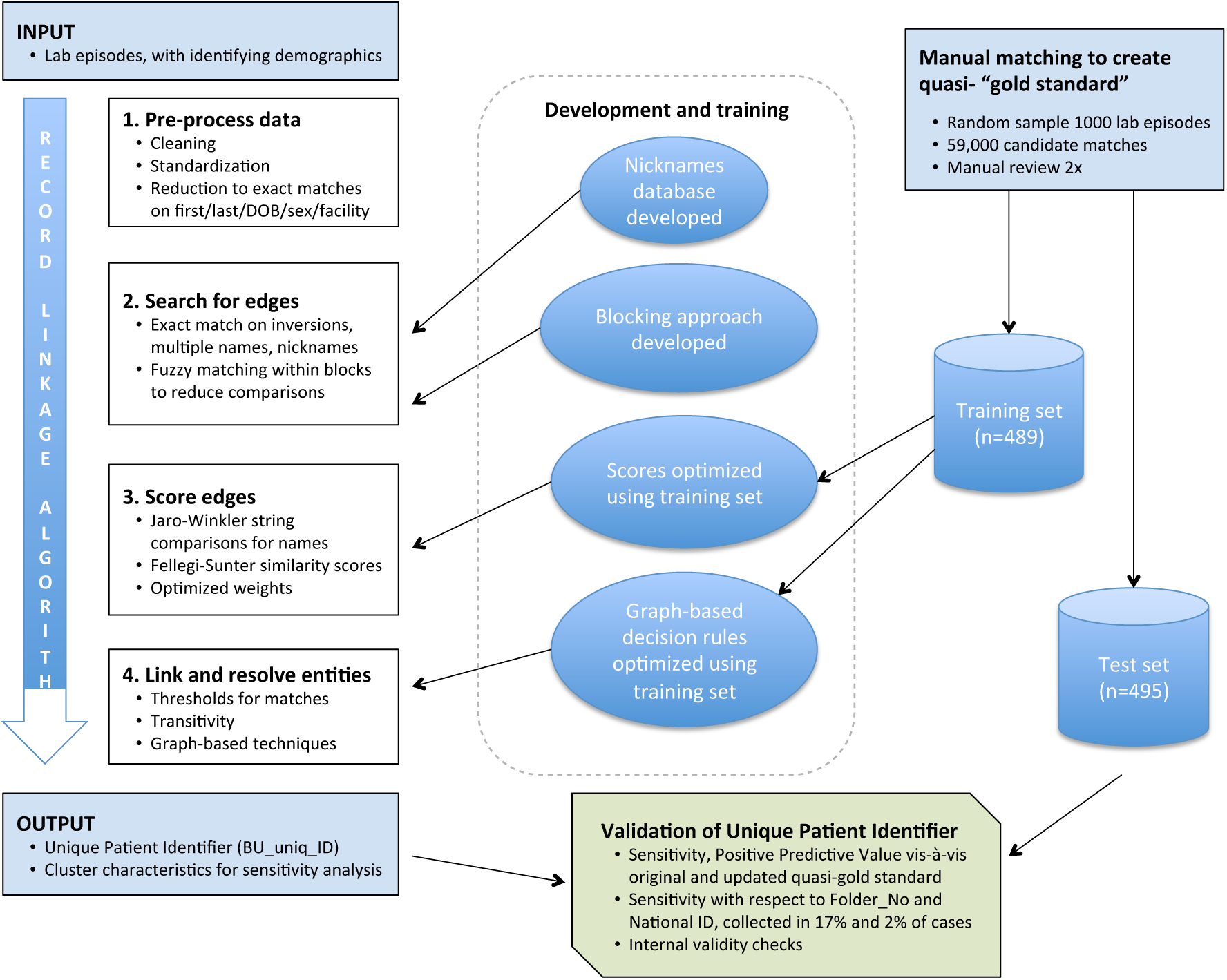
Graph-based probabilistic linkage schematic. Steps undertaken in linkage exercise.

### 3.2 Preprocessing

Pre-processing is a standard first step in record linkage. We began with the complete listing of Episode_No’s associated with CD4, VL, and other plausibly HIV-related tests in the NHLS CDW, and the available demographic information on these episodes. (Some Episode_No’s were associated with multiple tests if multiple tests were conducted on the same specimen. Eliminating these duplicate Episode_No’s reduced the number of rows from 154.8 million to 115.8 million).

In pre-processing the data, we had two goals. First, we sought to identify and exclude invalid laboratory results, e.g. those that were associated with a research study or routine quality control and thus did not reflect patients in the public-sector health system as well as specimens that had nonsensical identifying information due to a data entry or processing error and thus could not be linked. Second, for valid laboratory results, we sought to standardize the data fields, removing non-alphabetical characters, removing common prefixes (Mr, Ms), standardizing common last names, dropping redundant initials, and replacing as missing if the name did not exist, e.g. “No Name”, “Unknown”, etc. **Table 2** lists the pre-processing steps that we conducted.

**Table 2.**
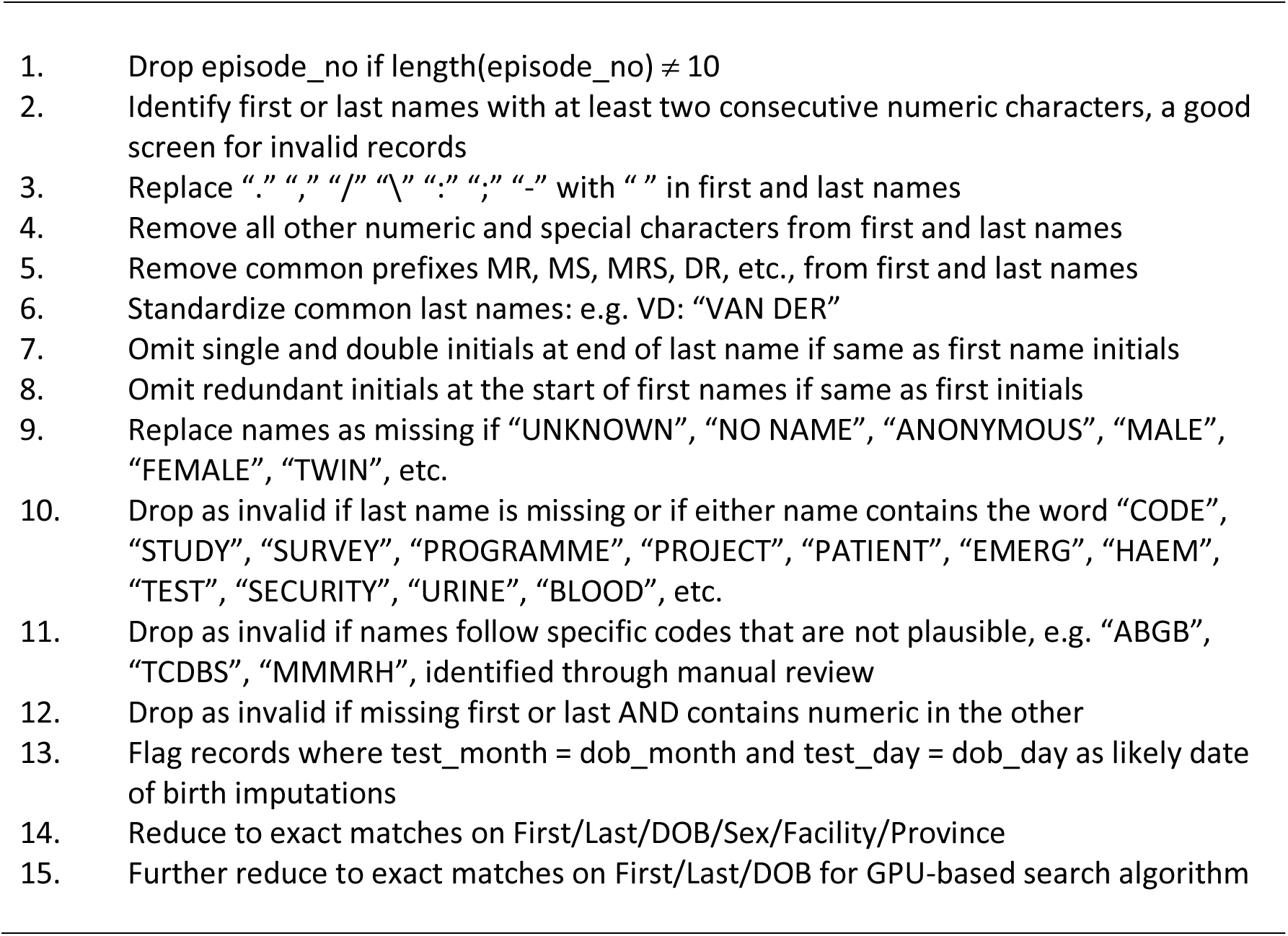
Preprocessing steps

After pre-processing and eliminating invalid results, we assessed the distributions of different first and last names, years of birth, genders, facilities, and provinces in the data, information that would be used in the scoring step below. Months and days of birth were assumed to be uniformly distributed within years, however we captured information on the distribution of year of birth to account for the non-uniform distribution of HIV patients. Each patient specimen was associated with a vector of probabilities based on the distribution in the full dataset, i.e. Pr(first = “John”), Pr(last = “Smith”), Pr(YOB = 1975), Pr(Gender = “M”), Pr(Facility = “ABCD”), Pr(Province = “EFG”).

To reduce the size of the dataset used in the linkage, we then collapsed the dataset to unique combinations of available identifying information – i.e., exact matches on first name / last name / date of birth / gender / facility / province. We created a crosswalk (linking file) from the specimen record identifier (Episode_No) to the exact matched identifier (EM_ID_plus). This reduced the size of the dataset from 115.8 million Episode_No’s to 62.8 million EM_ID_plus identifiers. These EM_ID_plus identifiers formed the nodes in the graph-based entity resolution step. To determine which nodes belong to the same patient, we searched for and scored edges between the nodes and then analyzed the resulting clusters.

### 3.3 Search

Our linkage was implemented on all unique sets of identifying information (n = 62.8 million). A complete n X n comparison of the dataset would require scoring 3.9 quadrillion comparisons, beyond our computing power. We therefore used several targeted and overlapping search strategies to reduce the number of comparisons. Our approach yielded about 433 million comparisons, reducing the number of required comparisons by a factor of 9 million.

Our primary search strategy was to assess all comparisons of the 63M cleaned, exact-matched, pre-processed records within an 11-year moving window on year of birth (**Table 3**). If the difference between two years of birth was 11 years or less, then we used the Jaro-Winkler algorithm to measure the similarity of first name pairs (first_JW) and last name pairs (last_JW). Jaro-Winkler similarity scores are on a 0 to 1 scale, with 1 representing an exact match. We compared all records within this year of birth window without further blocking (i.e. without requiring exact matches on other characteristics). In contrast to the common practice of blocking on initials, this approach allows for detection of similar names even when the initial letters differed (e.g. Carl ~ Karl). Records were retained if first_JW*0.6 + last_JW*0.4 exceeded 0.9. (The greater weight given to first name in this screening step was the result of initial investigations of the training data, which suggested that first name had more discriminating power than last name.) To execute this search strategy quickly, we developed a programme in C that could be run on parallel processors simultaneously (500 graphic processing units, or GPUs).

Though broad, the moving window search strategy could miss cases in which the difference in years of birth was greater than 11 years or where the name similarity was low. As a second search strategy, we supplemented the moving window with several targeted blocking approaches described in **Table 3**. For each of the following – first name, last name, DOB, and the combination of sex and facility – we allowed for fuzzy matching on that variable if there was an exact match on all other variables.

Third, we conducted deterministic searches for matches based on first/last name inversions, matches on multiple first or last names, and matches on a database of nicknames and common alternate names that we developed using statistically guided search with manual review. To develop the nicknames database, we identified all pairs of exact-matched ID’s in which the first name differed but the last name and date of birth were the same. We then counted the number of times a particular pair of first names occurred. A single occurrence could easily happen by chance, however multiple occurrences of the same pair of first names with different last names and dates of birth would suggest that the pair reflects a common nickname or misspelling. Restricting to first name pairs that appeared at least five times in the database, we constructed a list of 15,000 potential nicknames. Research staff at HE^2^RO fluent in the major South African languages then identified valid pairs from the list.

The result of these search strategies was a list of edges (pairs of nodes) that were of sufficient interest to be scored. The distribution of edges is displayed in **Table 3**.

**Table 3.** **Search strategy featuring multiple passes**

| Search Method | Number of Edges<br>(less duplicates) |
| --- | --- |
| 1. <u>Moving window</u> | 310,469,082 |
| Fuzzy match first and last if DOB within 11 years; retain if name similarity above threshold; run on GPUs |  |
| 2. <u>Overlapping blocks</u> |  |
| 2.1 Fuzzy match first if exact on all other variables | 10,586,682 |
| 2.2 Fuzzy match last if exact on all other variables | 6,840,290 |
| 2.3 Exact, except for DOB | 62,579,774 |
| 2.4 Exact, except for sex/facility | 32,413,676 |
| 2.5 Exact, except for first missing | 95,306 |
| 2.6 Exact, except for DOB missing | 3,349,716 |
| 3. <u>Deterministic comparisons</u> |  |
| 3.1 Name Inversions; exact DOB | 471,299 |
| 3.2 Multiple first name; exact last/DOB | 2,100,920 |
| 3.3 Multiple last name; exact first/DOB | 55,924 |
| 3.4 Nicknames; exact last/DOB | 3,959,943 |
| **TOTAL EDGES, LESS DUPLICATES** | **432,922,612** |

### 3.4. Scoring

We followed an adapted Fellegi-Sunter^6^ approach to score potential matches. Pairs of records were evaluated for similarity across each of six domains independently: first name, last name, date of birth, gender, province, and facility. Scores were assigned based on whether there was a match in each domain, and then the scores across the domains were aggregated using a weighted sum to determine a total similarity score. Comparing records *i* and *j*, the Fellegi-Sunter formula assigns scores for each domain *k* as follows:

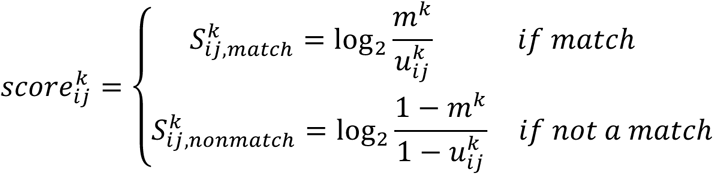

The “m-probability”, *m*^*k*^, is the probability of observing a match on domain *k* if the two results in fact belong to the same patient. The “u-probability”,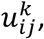 is the probability of observing a match if the two results in fact *do not* belong to the same patient, i.e. the false positive rate. The “m-probability” is a function of the data generating process for differences in identifying information being recorded for the same patient in a particular domain, including, e.g., typographical errors, as well as rates of transfer in the case of discordant facilities. The “m-probabilities” were estimated in the manually-matched training data and were assumed to be constant across the whole database (hence not indexed by i,j). (Similar estimates of the m-probabilities were obtained in the full database after implementing the algorithm.) The “u-probability” is a function of the frequency of the response values for record *i* and record *j* in domain *k.* The probability that another patient has exactly the same value for domain *k* can be estimated by the probability mass for that value in the database, e.g. Pr(gender=F), Pr(first=John). The less common a response value, the smaller the u-probability and the more credit given in the event of a match. When *i* and *j* differ, they have different u-probabilities. To avoid mistaking typographical errors for rare names, we defined 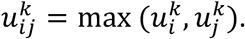 The values of the domain-specific scores, 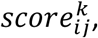 can be positive (if a match) or negative (if not a match). Missing data will yield a score of zero for that domain.

For gender, facility, and province, we scored pairs of records using the binary match/non-match designation built into the formula above. For first and last names, typographical and hearing errors can lead to slight differences, which are not clearly a match or non-match. Following Herzog et al (2007)^3^, we adapted the Fellegi-Sunter formula to account for fuzzy string matches using the Jaro-Winkler scoring algorithm for string comparisons.^28,29^ The Jaro-Winkler similarity metric is based on the share of characters in each string that also occur in a similar location in the other string. Additional weight is given to strings that match on initial letters. The similarity score is scaled from 0 to 1. The Jaro-Winkler similarity score has been shown to perform as well or better than other string comparison metrics.^3,30^ For nicknames, first/last inversions, and matches on multiple middle names, we replaced the Jaro-Winkler score with 0.95, to account for the small decrement from an exact match. For first and last names, 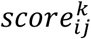 was defined as:

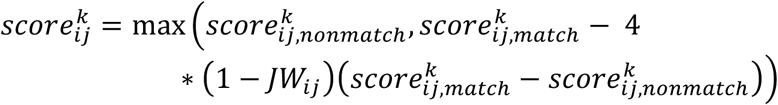

where 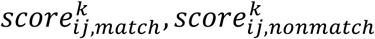 are the scores if a match and if not a match, respectively, and *γ* is the Jaro-Winkler similarity score. The formula assigns the non-match score if *JW*_*ij*_ ≤ 0.75 and linearly interpolates between the non-match and match scores of the Jaro-Winkler similarity is between 0.75 < *JW*_*ij*_ ≤ 1.

The 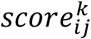 for date of birth was based on the Fellegi-Sunter formula, but also incorporated additional information about the data generating process giving rise to variation in recorded dates of birth. In particular, when patients provided an age rather than an exact date of birth, then the year of birth in the CDW database was imputed by subtracting the age from the current year, and the month and day of birth in the CDW database were imputed using the month and day of the laboratory test. Therefore, when the month and day of the laboratory test were identical to the month and day of birth, we assumed that the month and day were in fact missing 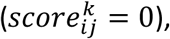 and the year of birth was assumed to be imprecisely reported, giving partial credit to matches with close but not identical years of birth.

The Fellegi-Sunter formula scores matches based on the amount of information they contain. Therefore, an exact match on date of birth or first/last name would be worth substantially more than an exact match on province or gender, which are more likely to occur by chance. **Appendix Figure 1** shows the distribution of scores for the different elements among “true” matches in the training set. In general, matches on dates of birth and names contributed the most to similarity scores.

After computing a 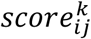 for each of the six components of the comparison vector, a total similarity score was calculated as a weighted sum:

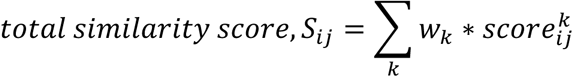

In choosing values of the weights, *w*_*k*_ the goal is to maximize the discriminating power of the total similarity score *S*_*ij*_ to distinguish true matches from true non-matches. **Figure 2** shows the distributions of total similarity scores among true matches and true non-matches in the manually-matched training data. The optimal weights *W*_*k*_ are those that separate these distributions as much as possible, giving high scores to true matches and low scores to true non-matches. We used the manually-matched training data to optimize *w*_*k*_ as follows:

**Figure 2.**
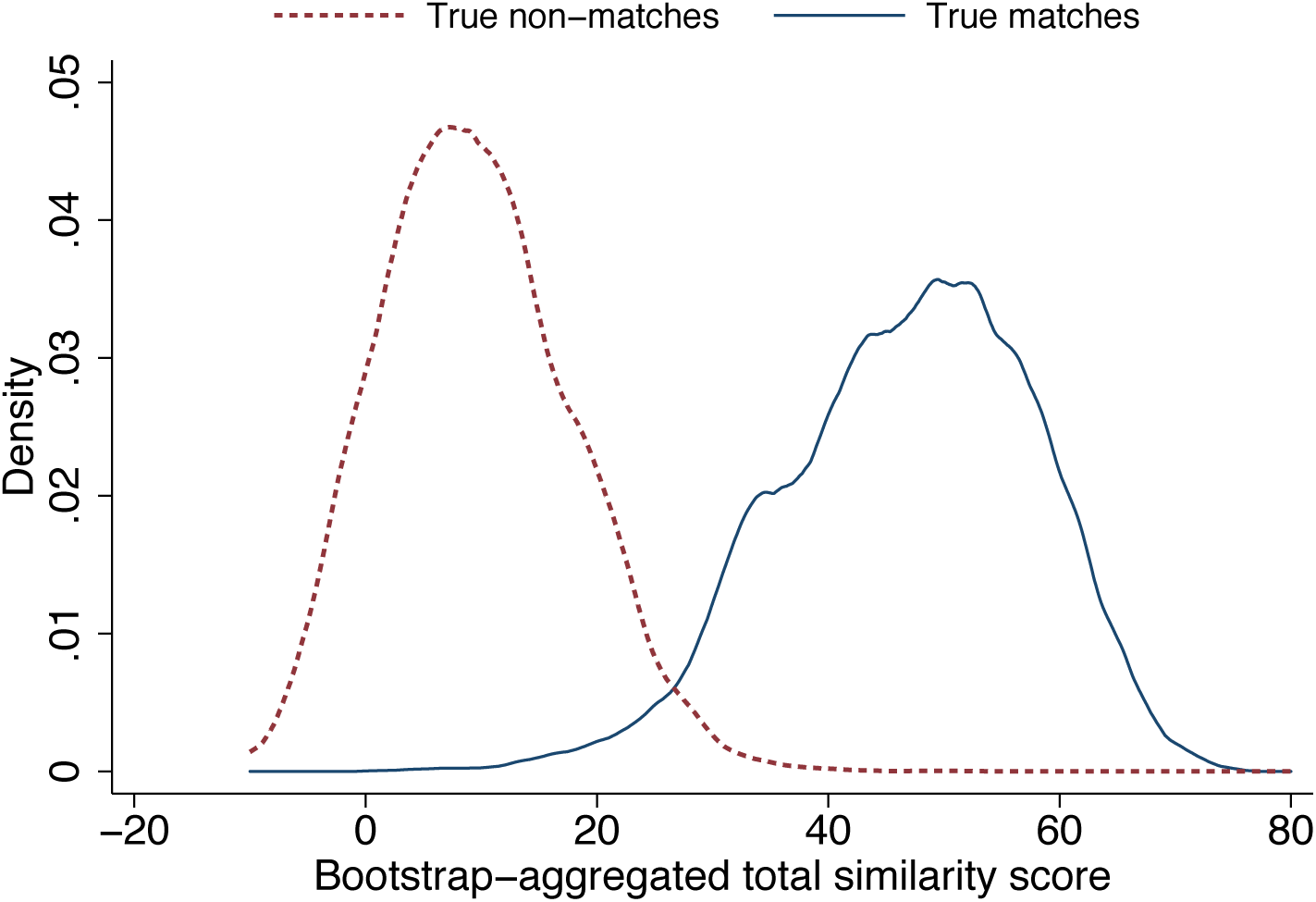
Distribution of scores in training data. Figure shows the distribution of bootstrap-aggregated “total similarity scores” for true matches and true non-matches. The bootstrap-aggregation procedure is described in the text below.

To optimize the weights, we defined an objective function and then optimized it using R’s `optim` package. Consider the total similarity score *S* = *S(w)* which is a function of the weights to be optimized. One way to assess the discriminating power of *S* is to propose a threshold decision rule 1[*S* > *τ*] (i.e., 1 if *S* > *τ*, 0 if *S* ≤ *τ*), which considers a comparison to be a match if the value of the similarity score is above some threshold *τ* (which could be denoted as a vertical line on **Figure 2**). For each candidate scoring function *S* and some value of *τ*, PPV and Sensitivity can be evaluated. Because the graph-guided entity resolution step allows the matching threshold to vary based on the density of the network, we were interested in discriminating power across the whole range of possible values of τ, not just at a single optimum. We computed *PPV*(*τ*|*S*) and *Sen*(*τ*|*S*) for the set of indicator functions 1[*S* > *τ*] across the range of thresholds τ supported by the training data. **Figure 3** shows how *PPV*(*τ*|*S*) varies with *Sen*(*τ*|*S*), with different values of *τ* tracing out the curve. At higher values of *τ*, the indicator function 1[*S* > *τ*] would have high PPV but low Sensitivity, as most true matches have scores less than *τ*. At lower values of *τ*, Sensitivity increases but PPV falls. Thus, as *τ* moves left to right in **Figure 2**, the line in **Figure 3** moves from bottom-right to top-left. We defined our objective function as the area under the Sensitivity-PPV curve. Because Sensitivity is defined with respect to the true matches, this AUC_Sen-PPV_ metric can be interpreted as the average PPV across percentiles of the true matches.

**Figure 3.**
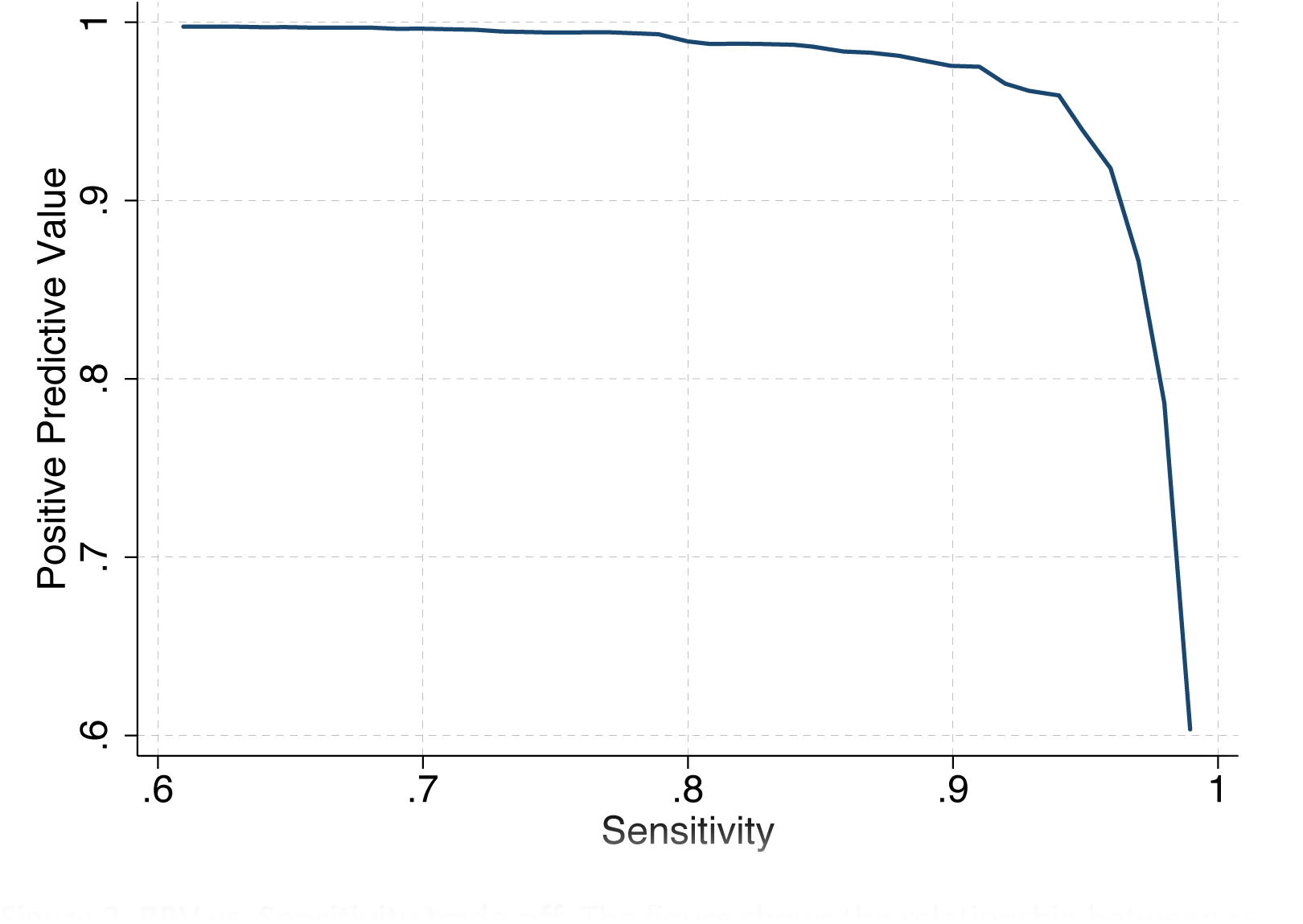
PPV vs. Sensitivity trade-off. The figure shows the relationship between sensitivity and specificity based on a single-threshold decision rule. Weights in the total similarity score were chosen via bootstrap aggregation to maximize area under the curve.

We created an R function that took as an input a hypothesized set of weights *w*_*k*_, and then computed the total similarity score *S* using those weights, and calculated the value of the objective function *AUC*_*Sen-PPV*_ in the training data. We chose the weights that maximized *AUC*_*Sen-PPV*_ in the training data using R’s `optim` package. In order to avoid over-fitting the weights to the training data, we applied *bootstrap aggregation or “bagging”^31,32^* to this optimization procedure. The above procedure was repeated for five hundred bootstrapped samples from the training data, generating 500 sets of optimal weights. We inspected univariate and bivariate distributions of the optimal weights across the 500 bootstrapped samples to assess stability. We found no evidence of multiple optima. We then calculated the simple average across the 500 sets of weights, resulting in a final set of weights, which we applied to the vector of domain scores to create a total **bootstrap-aggregated similarity score**, *S*_*BAG*_, shown in **Figure 2**, which predicts pairwise matches without overfitting the training data. The final weights were:

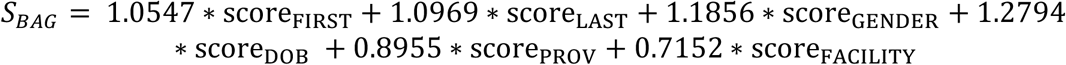

The bootstrap-aggregated similarity scores are on an arbitrary scale, ranging from about −10 to 80. To facilitate interpretability, we transformed *S*_*BAG*_ into “true match” probabilities by running a logistic regression model of the manually-matched training data (1=match, 0=not a match) on *S*_*BAG*_ and using the coefficients to obtain predicted probabilities. **Figure 4** shows the fit of this model and illustrates the range of values of *S*_*BAG*_ for which the match is uncertain.

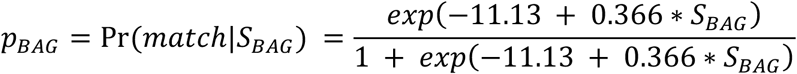

**Figure 4.**
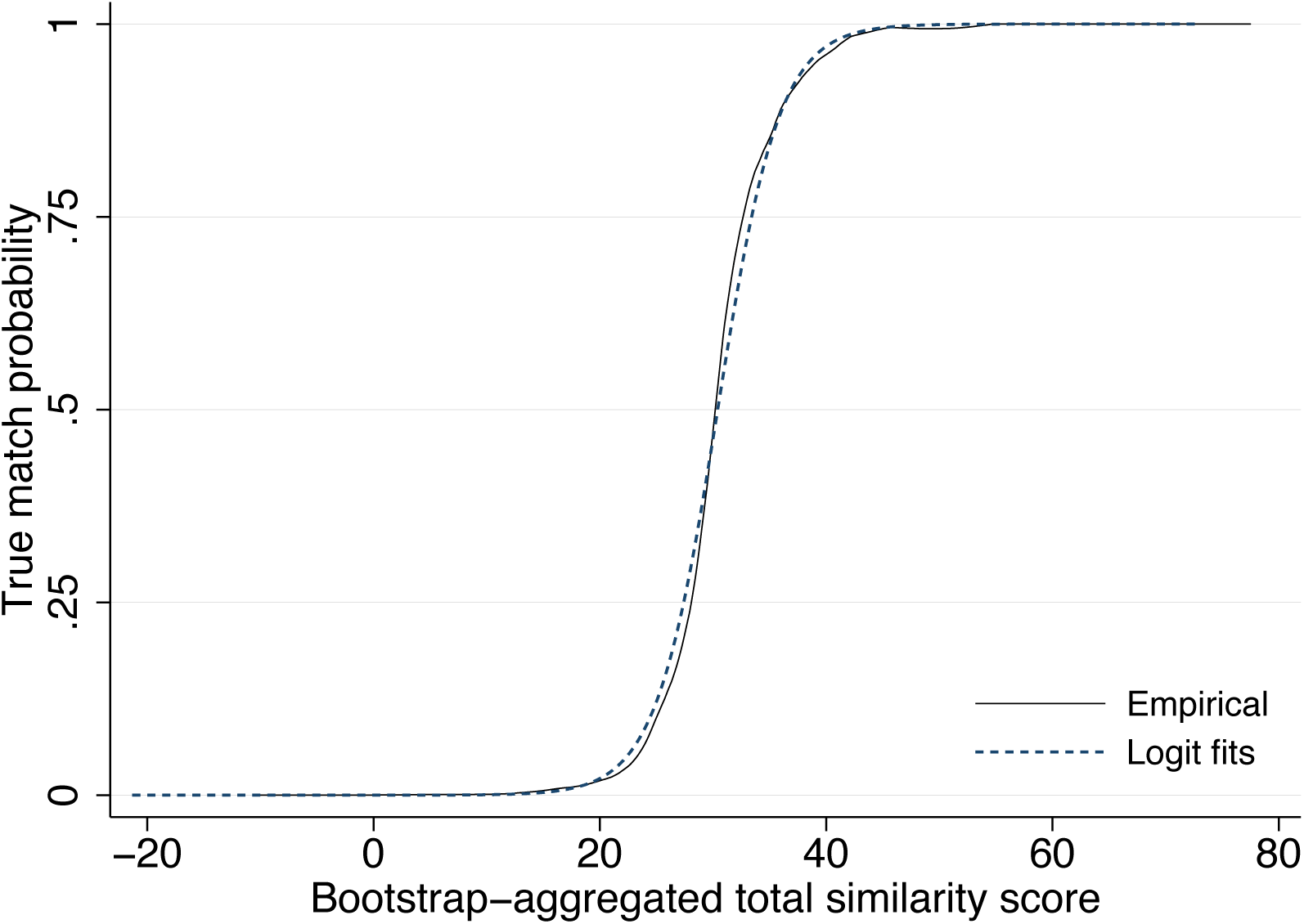
Estimating match probabilities. Figure illustrates parameters for a logit transformation of the similarity scores into predicted probabilities based on the training data.

Finally, in the graph-based entity resolution step that follows, edge weights are specified as a distance (rather than similarity) metric, with higher values reflecting more dissimilar records and greater distance in the network. We therefore define edge weights as:

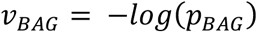

### 3.5 Graph-based entity resolution

Our initial plan was to choose the single best threshold value for *p*_*BAG*_ across the whole dataset that would minimize over-and under-matching errors. However, after implementing this strategy, we found that choosing a single threshold led to substantial over-matching, substantial under-matching, or both. Because of the large size of the dataset, thresholds that were low enough to achieve desired sensitivity also linked together records that should not have been linked. As a result, our initial efforts led to very large clusters of observations (e.g., >1M records linked together as one patient). The previous record linkage effort by CDW also encountered this issue.

To solve this problem, we used graph concepts to guide the identification of unique patients. The scored comparisons can be thought of as weighted edges in a network, where the nodes represent unique sets of identifying information. Using R’s igraph package, we formed the full graph (network) defined by these nodes, edges, and assigned edge weights (*v*_*BAG*_) to the edges. Our goal was to identify clusters that represented unique patients. We developed our approach based on the following logic:

1. A cluster of nodes cannot reflect just one single patient if the two most dissimilar nodes in the cluster do not belong to the same patient
2. A cluster of nodes *likely* reflects a single patient if the two most dissimilar nodes in the cluster belong to the same patient
3. The shortest-weighted-path distance between the two furthest nodes in the cluster is the weighted diameter, which is the sum of the edge-specific weights.
4. Distance reflects dissimilarity, and weighted diameter thus captures the dissimilarity of the most dissimilar nodes in the cluster.

By this logic, the weighted diameter can be interpreted as a measure of plausibility for whether the cluster represents a single patient. Because we defined the edge weights as *v*_*BAG*_ = − *log p*_*BAG*_, the sum of the edge weights along the shortest weighted path between the furthest nodes is equal to the log-product of probabilities along this path. The weighted diameter of cluster *c* is:

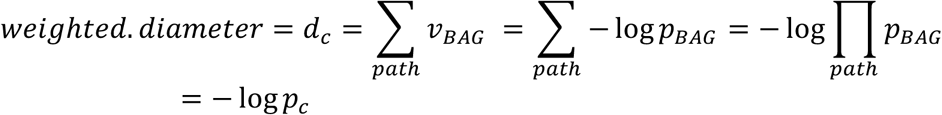

where *p*_*c*_ = exp (−*d*_*c*_) is defined as ∏_*path*_ *p*_*BAG*_ and reflects the similarity between the two farthest nodes. In the special case in which the weights for each of the edges along the diameter path are independent, then *p*_*c*_ is interpretable as the probability that the two most dissimilar nodes belong to the same patient. Independence would arise if the database errors leading to edges along the diameter path were orthogonal. For example, suppose A and B differ because of a typographic data entry error in the last name of A; and B and C differ because C reported age in lieu of date of birth. It is likely these errors are independent. (We note, however, that it is easy to construct cases of positive or negative dependence.) For a two-node cluster, the weighted diameter is simply the single edge score.

An attractive feature of weighted diameter is that it is a relevant metric regardless of the size of a cluster. A two-node cluster with a single edge can be subjected to the same decision rule as a 10-node cluster with a diameter that traverses four nodes. In both cases, weighted diameter captures how likely it is the two most dissimilar nodes in the cluster belong to the same patient. And in both cases, we set a minimum similarity threshold value of *p*_*c*_ ≥ *θ*;, which corresponds to a maximum distance threshold value of *d*_*c*_ ≤ − log *θ* for the weighted diameter of each cluster. Clusters with a weighted diameter less than the − log *θ* threshold were deemed plausible patients and moved to the final dataset. If clusters had a weighted diameter greater than that threshold, then the lowest scoring edges were deleted and the cluster was reassessed. For clusters with >10 edges, we deleted the lowest-scoring 10% of edges; for clusters with <10 edges, we deleted the single lowest-scoring edge. This process was repeated iteratively until all clusters had a weighted diameter less than − log *θ* and had been moved to the final dataset. (See **Figure 5** for an illustration of how a large cluster was broken up.) The final dataset consisted of a graph of the complete database in which all clusters were deemed plausible patients. We labeled the cluster identifiers as the BU_uniq_ID (Boston University unique identifier) and exported a file assigning all nodes to BU_uniq_ID’s.

**Figure 5.**
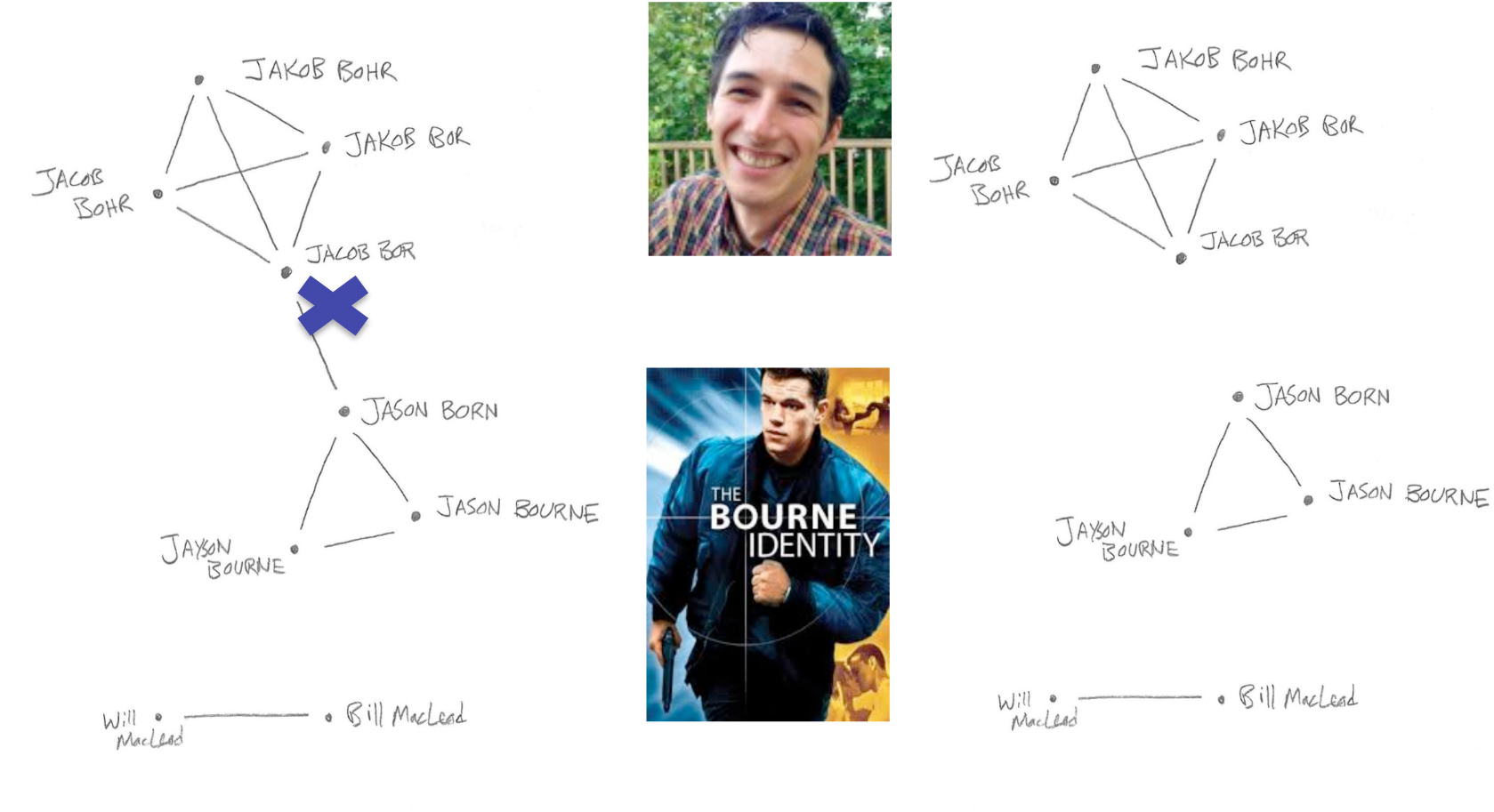
Graph example (to be updated with real example) Figure shows how graph-based entity resolution breaks up a large cluster that resulted from the initial stage of probabilistic matching. Names are changed to preserve privacy.

Mapping large graphs is computationally intensive. To speed up the approach, we partitioned the graph of the full dataset into discrete sets of clusters and ran the graph-based entity resolution code separately on these partitions. Additionally, at the outset, we restricted clusters to no more than 100 nodes, dropping low-scoring edges until all clusters met this criterion. (We considered it implausible a patient would have more than 100 sets of identifying information.)

To choose the optimal threshold for the weighted diameter, we conducted a grid-search over a range of possible values for the threshold. We assessed thresholds of *θ* = 0.25,0.3,… 0.95. For each threshold, we implemented the graph-based entity resolution step for the full database, creating a different version of the BU_uniq_ID. We then assessed performance of these BU_uniq_ID’s with respect to the training data in terms of Sensitivity and PPV.

**Figure 6** shows the results of this grid search, plotting the Sensitivity and PPV of the algorithm when using different thresholds. The higher the value of *θ*, the more clusters are broken up and the lower the Sensitivity, but higher the PPV. The less the clusters are broken up, the higher the Sensitivity, but the lower the PPV. The question of what the threshold should be depends on how the user values Sensitivity vs. PPV. Higher thresholds will lead to lower Sensitivity and higher PPV, and vice-versa. In the extreme, a threshold of 1 will be equivalent to exact matching. One common approach is to maximize the F-measure, which is the harmonic mean of Sensitivity and PPV: *F* =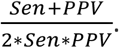 In **Figure 6**, shaded bands reflect isoquants (contours) for the F-measure.

**Figure 6.**
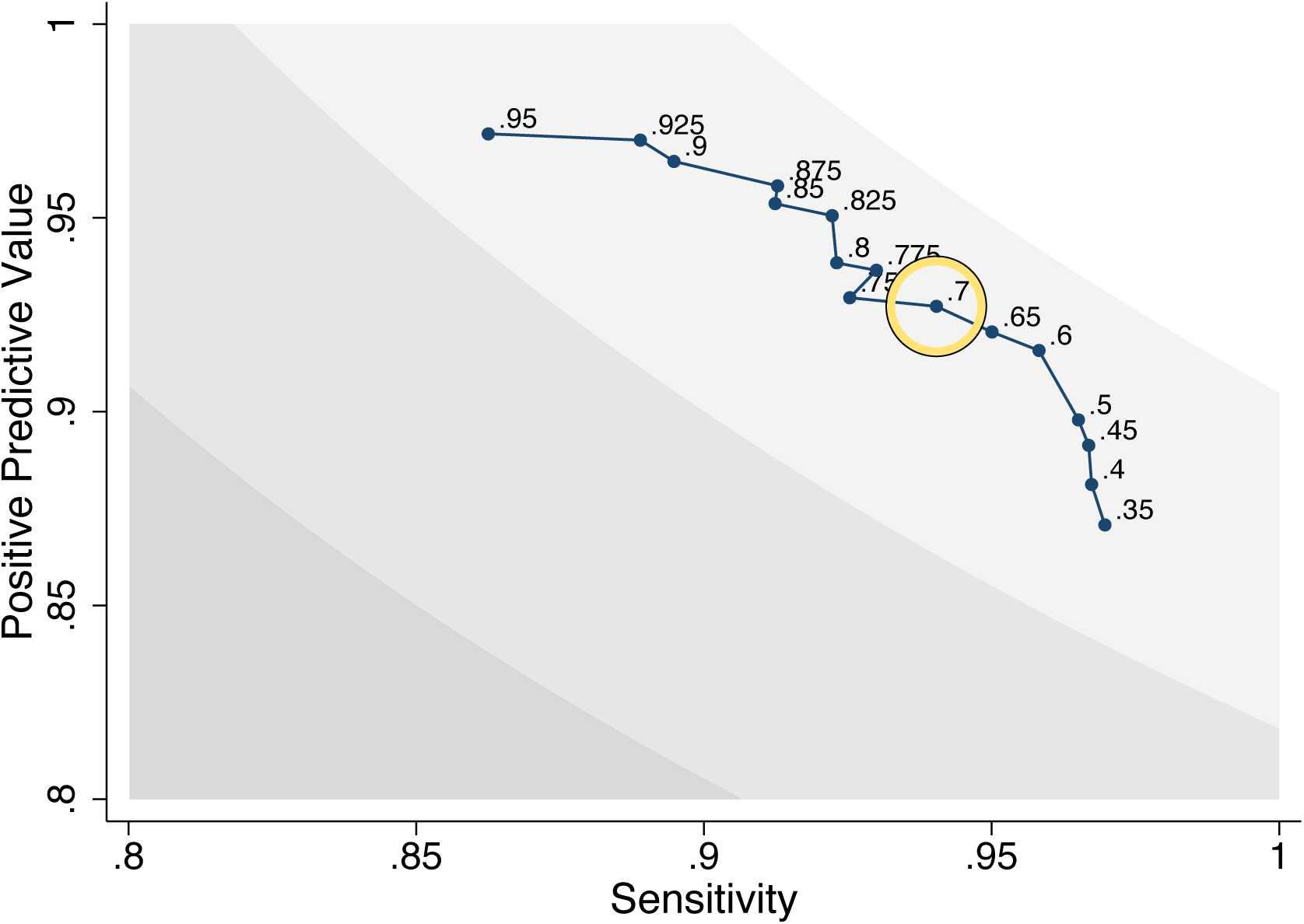
Optimizing graph-based entity resolution. Figure shows Sensitivity and PPV of the linkage algorithm in the training data, using different thresholds for the weighted diameter.

Based on the grid search we decided to use a weighted diameter threshold of *θ* = 0.7. Although 0.7 was outperformed in the training data by 0.825 and 0.6, we considered that the underperformance of 0.7 was likely a reflection of noise in the training data.

### 3.6. Computational performance

The graph-based record linkage pipeline was implemented on Boston University’s Shared Computing Cluster (SCC) in a secure environment. Most tasks were implemented in Stata-MP 15.0. The moving window search strategy was written in C/CUDA and executed on the cluster using graphic processing units (GPUs). The optimization of weights via BAGging and the graph-based entity resolution step were conducted in R 3.2.3. **Table 4** shows computational time of each step. During training, Stata was used to assess performance of the algorithm compared to manually-coded data.

**Table 4.** **Computing procedures and time**

| Linkage Step | Software | Computing Time |
| --- | --- | --- |
| 1. Pre-processing | Stata-MP | ~3 h |
| 2. Search | C/CUDA, Stata-MP | ~5 h |
| 3. Scoring | R, Stata-MP | ~3 h |
| 4. Graph-based entity resolution | R | ~4 h |

Notes: Steps 1,3 and 4 were executed using Intel Xeon Processor E5-2650x. Step 2 was executed using 2 NVIDIA's V100 GPUs with 16GB of memory.

## 4. LINKAGE RESULTS AND VALIDATION

### 4.0 Preliminary results of the linkage project

Results of an earlier version of the linkage were presented in April 2016, based on data on all CD4 counts and viral loads collected January 2004 – first quarter of 2015. Comparison of the results to the manually matched training data revealed estimated Sensitivity of 91.0% and PPV of 90.5%. The results presented here reflect updated data, improvements to the algorithm, and further review of the manually-matched data.

### 4.1 Results of the graph-based record linkage

The database included all CD4, VL, HB, ALT, CrAg, CrCl, and HIVPCR records in the NHLS CDW from January 2004 – December 2016. (An additional 978 results were inadvertently included from outside this range.) We started with 239.8 million CD4 counts and viral loads, associated with 117.5 million specimens (**Figure 7**). After preprocessing and removing exact duplicates, we were left with 62.8 million unique sets of identifying information. Our algorithm identified 11,632,222 unique patients from these data, who had at least one CD4 count or VL. These 11.6 million HIV patients had 70.9 million specimens corresponding to 97.7 million CD4, VL, or one of the other laboratory tests used in HIV monitoring: HB, ALT, CrAg, CrCl, PCR; 44.7 million specimens were excluded because they corresponded to one of the other laboratory tests but were not linked to a patient with a CD4 or VL. **Appendix Table 1** displays the distribution of tests by type and year in the dataset.

**Figure 7.**
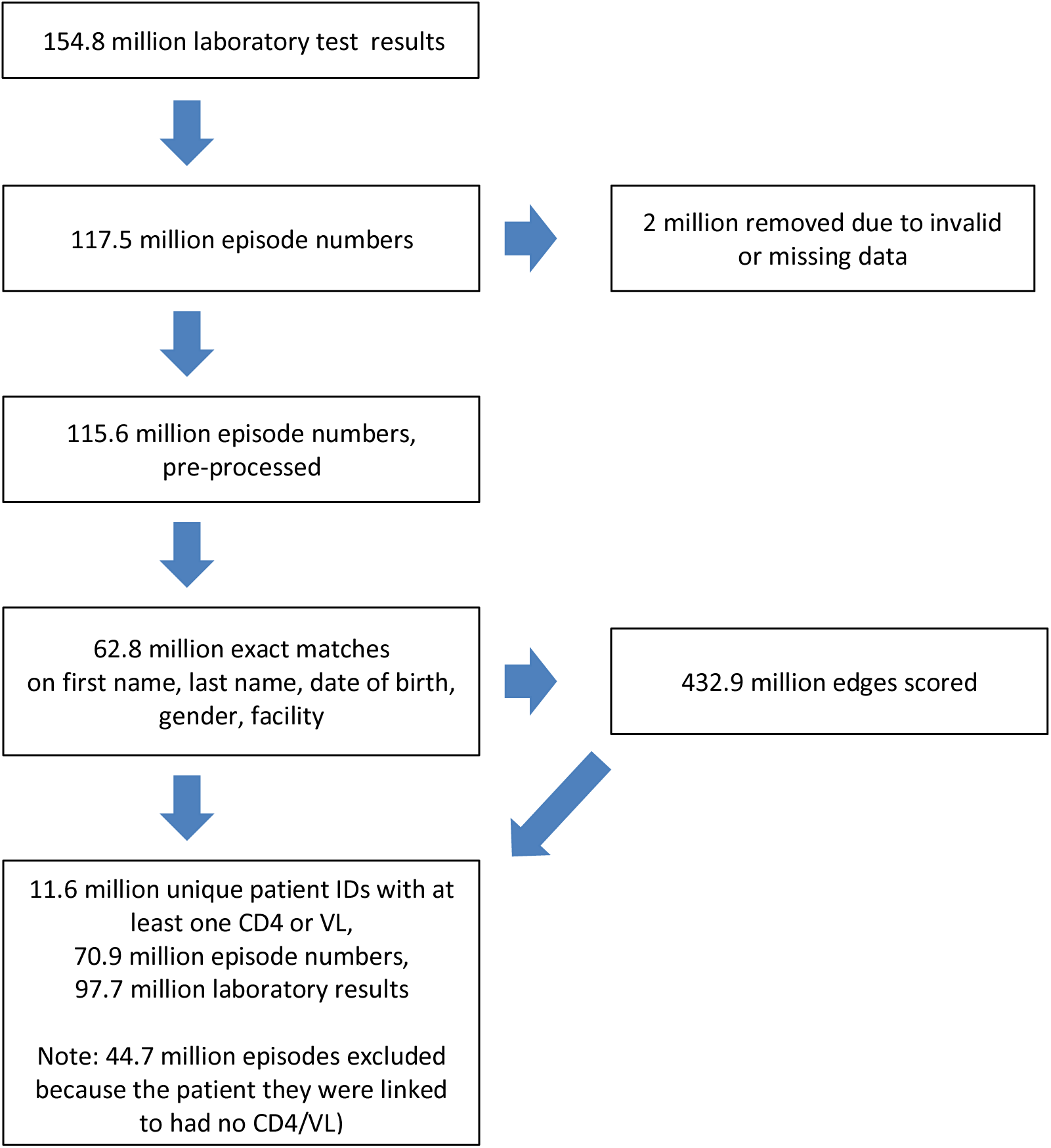
**Results of graph-based record linkage: flowchart**.

**Table 5a** shows the distribution of numbers of nodes in each cluster associated with an individual patient. Nearly half of patients had multiple sets of identifying information. However, just one percent had 10 or more sets of identifying information. The maximum was 71. **Table 5b** displays the distribution of laboratory episodes per patient. The median number of episodes was 3 with an IQR of 1 to 8. 28% of patients had just a single specimen. About half of patients had between 2 and 7 results. And a quarter had 8 or more specimens. Only 1% of patients had more than 38 specimens. **Table 5c** displays the distribution of laboratory tests across test type in the linked cohort. Note that there can be multiple laboratory tests per specimen. There were 20.1 million viral loads, 32.5 million CD4 counts, and 45.0 million other tests conducted 2004-2016 in these patients.

**Table 5a.**
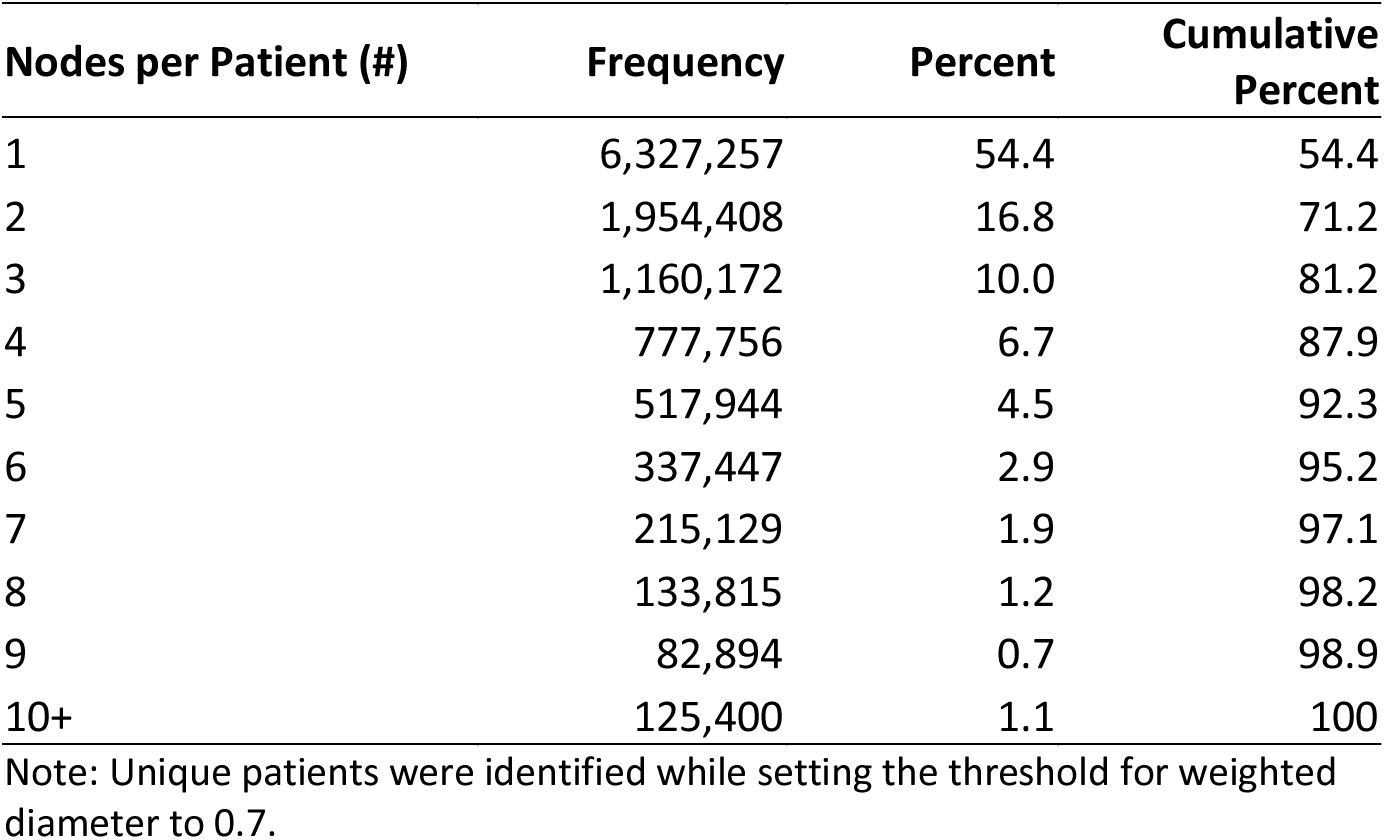
**Number of sets of identifying information (nodes) per BU_uniq_ID**

**Table 5b.** **Number of specimens per patient (wd=70)**

| **Episodes per patient (#)** | **Frequency** | **Percent** | **Cumulative Percent** |
| --- | --- | --- | --- |
| 1 | 3,254,841 | 28.0 | 28.0 |
| 2 | 1,719,683 | 14.8 | 42.8 |
| 3 | 1,136,989 | 9.8 | 52.5 |
| 4 | 861,037 | 7.4 | 59.9 |
| 5 | 668,485 | 5.8 | 65.7 |
| 6 | 552,676 | 4.8 | 70.4 |
| 7 | 463,988 | 4.0 | 74.4 |
| 8 | 392,856 | 3.4 | 77.8 |
| 9 | 333,329 | 2.9 | 80.7 |
| 10-14 | 1,060,424 | 9.1 | 89.8 |
| 15-19 | 503,270 | 4.3 | 94.1 |
| 20-49 | 648,691 | 5.6 | 99.7 |
| 50-99 | 34,058 | 0.3 | 100.0 |
| 100+ | 1,895 | 0.0 | 100.0 |
| Total | 11,632,222 | 100.0 |  |

Note: Unique patients were identified while setting the threshold for weighted diameter to 0.7.

**Table 5c.** **Laboratory tests among HIV patients identified in the linked cohort**

|  | Test type | Number of Results | Percent |
| --- | --- | --- | --- |
| *HIV monitoring labs* | CD4 | 32,541,453 | 33.3 |
|  | Viral Load | 20,152,341 | 20.6 |
| *Labs used in ART work-up and HIV confirmatory testing* | ALT | 14,510,715 | 14.9 |
|  | CRAg | 777,627 | 0.8 |
|  | Creatinine | 5,558,418 | 5.7 |
|  | HIV Elisa/PCR | 3,261,202 | 3.3 |
|  | Hemoglobin | 20,907,431 | 21.4 |
|  | **Total** | **97,709,187** | **100** |

Note: Unique patients were identified while setting the threshold for weighted diameter to 0.7.

### 4.2 Validation of the linkage

We validated the linkage algorithm using three approaches. Our primary approach was to assess the Sensitivity and PPV of the BU_uniq_ID compared to the manually-matched validation set, based on our random sample of CD4/VL results from Fall 2014. We also computed corrected Sensitivity/PPV measures after a second round of manual review of this validation set. We computed standard errors/confidence intervals using the cluster bootstrap, resampling reference Episode_No’s (specimens) from the validation set.

We also assessed sensitivity in relation to two other identifiers available for a portion of the database: Folder Number (17% of episodes) and National ID Number (2% of episodes). We cleaned folder numbers eliminating garbage codes, e.g. “NO FOLDER NUMBER”, codes with fewer than 7 digits (which could correspond to multiple patients), and other codes that were likely to be non-unique or errors. Manual inspection revealed that even after cleaning, there were a substantial number of folder numbers that were clearly non-unique (including some associated with over 10 patients). However, we could not detect systematic patterns to enable further cleaning. Sensitivity estimates based on the folder number are likely to be biased downwards due to the presence of these non-unique codes. To ameliorate this problem, we also combined folder numbers with facility identifiers because different facilities may use the same folder number (however, in doing so we also eliminate transfers). National ID numbers were restricted to valid ID numbers, defined as those containing exactly thirteen digits and for which the “check digit” value in the final number was valid based on the Luhn algorithm.

Third, we assessed various metrics for each identifier (BU_uniq_ID_wd70, EM_ID, CDW_uniq_ID) in the dataset such as the number of unique patients, numbers of patients with many specimens, numbers jumping back and forth across provinces, numbers with multiple sexes, numbers with large CD4 count swings (>500 cells per year over a period of at least 6 months), and numbers with the first record being a viral load (by guidelines first test should be CD4).

Table 6, Appendix Table 2, and Table 7 display validation results. BU_uniq_ID (with a weighted diameter of 0.7) attained Sensitivity of 93.7% (95% CI 92 to 96) and PPV of 98.6% (95% CI 98 to 100) vis-à-vis the revised validation set. Results were broadly similar comparing BU_uniq_ID_wd70 to the original validation set and the training data.

**Table 6.** **Validity of Unique Patient Identifiers**

| Unique identifier | Sensitivity |  | Positive Predictive Value |  | F-measure |
| --- | --- | --- | --- | --- | --- |
|  | *Original* | *Corrected* | *Original* | *Corrected* | *Corrected* |
| **Exact match ID** | 59.5% | 53.4% | 100.0% | 100.0% | 69.6% |
| **CDW Unique ID** | 71.0% | 64.0% | 99.3% | 100.0% | 78.1% |
| **BU Unique ID (wd=50)** | 96.5% | 96.5% | 91.5% | 97.7% | 97.1% |
| **BU Unique ID (wd=70)** | 95.2% | 93.7% | 93.3% | 98.6% | 96.1% |
| **BU Unique ID (wd=90)** | 91.9% | 88.4% | 94.8% | 99.5% | 93.6% |

**Table 7.** **Additional validation results**

| Algorithm | Exact match | CDW unique ID | BU unique ID |
| --- | --- | --- | --- |
| *A. Numbers of patients identified* |  |  |  |
| Unique Patients | 20,212,961 | 18,459,757 | 11,632,222 |
| Ever on ART | 9,884,397 | 9,069,305 | 5,945,339 |
| On ART in 2015/16 | 5,515,572 | 5,280,204 | 4,191,525 |
| *B. Performance relative to manually-matched quasi-gold standard* |  |  |  |
| Sensitivity | 59.5% | 71.0% | 95.2% |
| Positive Predictive Value | 100.0% | 99.3% | 93.3% |
| Corrected Sen<sup>a</sup> | 53.4% | 64.0% | 93.7% |
| Corrected PPV<sup>a</sup> | 100.0% | 100.0% | 98.6% |
| *C. Performance relative to existing IDs* |  |  |  |
| Sensitivity relative to Folder_No (17% of specimens)<sup>b</sup> | 76.3% | 79.6% | 89.8% |
| Sensitivity relative to Folder_No-X-Facility<sup>b</sup> | 81.3% | 84.9% | 92.8% |
| Sensitivity relative to National ID (2% of specimens)<sup>b</sup> | 87.9% | 95.3% | 98.5% |
| *D. Measures internal to the dataset* |  |  |  |
| *Evidence of over-matching* |  |  |  |
| # with >10 nodes* | 6900 | 10,001 | 125,400 |
| # with >25 nodes* | 1110 | 207 | 206 |
| # with >100 specimens* | 1379 | 1303 | 1895 |
| # with >1000 specimens* | 0 | 3 | 0 |
| # with both M and F sex | 2.1% | 2.7% | 4.8% |
| # changing provinces >5X | 0.8% | 1.0% | 2.4% |
| # changing facilities >5X | 5.8% | 7.4% | 18.2% |
| # with ΔCD4 > 500 cells/year | 4.2% | 4.3% | 4.8% |
| *Evidence of under-matching* |  |  |  |
| # with VL and no prior CD4 | 13.4% | 13.2% | 6.5% |

\*Notes: Exact matches are on first, last, DOB. <sup>a</sup>Corrected sensitivity and PPV are based on re-review of the gold standard dataset. <sup>b</sup>Sensitivity, for Validation Approach 2, is defined as the proportion of links identified by folder number or national ID also identified by a given algorithm. Ever on ART is defined as ever having had a VL. On ART in 2016 is defined as any VL in 2015/2016, with a 2-year window to ensure that individuals who are substantially late for their 12-monthly VL are included. Nodes = unique sets of identifying information defined by first/last/DOB/gender/facility/province. Specimens = Episode\_No's.

The BU_uniq_ID achieved similar PPV as the existing CDW_uniq_ID and exact matching (98.6 vs. 100.0% vs. 100.0%), while attaining large improvements in Sensitivity (93.7% vs. 64.0% vs. 53.4%). Due to the greater sensitivity of the graph-based algorithm, the BU_uniq_ID identified smaller total numbers of patients relative to the CDW_uniq_ID and exact matching (11.6M vs. 18.5M vs. 20.1M) and fewer patients currently on ART (4.2M vs. 5.3M vs. 5.5M). 11.6M is at the upper range of plausible values for the total number of patients that have ever sought care in South Africa’s national HIV program. Indeed, as the Sensitivity estimates indicate, there is still scope for further matching – although our algorithm was not able to accurately identify further matches. Assessment of the distribution of cluster sizes shows that the BU_uniq_ID was able to increase sensitivity by substantially increasing the number of mid-to-large clusters (10-25 sets of unique identifying information) while having no effect on the number of very large clusters (>25 sets of unique identifying information). In fact, whereas the CDW_uniq_ID identified three “patients” with over 1000 specimens; the BU_uniq_ID identified no such patients.

Sensitivity of the BU_uniq_ID was very high (98.5%) vis-à-vis national ID numbers, albeit in the 2% of episodes that contained national ID numbers. Sensitivity was relatively high compared to folder numbers (89.8%) and when combining folder numbers with facility identifiers (92.8%). As a final indicator of improved sensitivity, the BU_uniq_ID cut in half the number of “patients” whose first test was a viral load (contrary to guidelines), from over 13% of “patients” identified by the CDW_uniq_ID or exact matching to 6.5%.

## 5. CONCLUSION

We developed and validated a record linkage algorithm that combined traditional scoring methods with graph-based concepts to guide the linkage. The graph-concept utilized – weighted diameter – captures the similarity of the most dissimilar nodes in a cluster and can therefore be used to identify clusters that could not plausibly reflect individual patients. Although our approach is not the first to use information on graph structure in record linkage, to our knowledge it is among the first to demonstrate the benefits of using information on weighted diameter in a large health dataset. Our approach incorporates information about both the size/shape of clusters and their locations within the broader network which is not traditionally utilized in record linkage procedures. Exploiting graph information has the potential to substantially improve the scalability of record linkage procedures in large datasets.

We applied the algorithm to the complete laboratory records from South Africa’s national HIV program, as compiled in the NHLS CDW. We identified 11.6 million unique patients with 97.7 million laboratory tests, from 61 million different sets of identifying information. Comparing the results to a manually-matched validation set, we achieved 93.7% Sensitivity and 98.6% PPV. We identified numbers of patients on HIV treatment similar to the numbers reported by South Africa’s NDOH.

By applying a novel graph-based record linkage algorithm to the NHLS database, we generated and validated a unique patient identifier, enabling longitudinal patient-level analysis and incorporation of longitudinal concepts (such as retention) into monitoring and evaluation dashboards. To our knowledge, the linked NHLS database represents the first nationwide HIV cohort in any low-or middle-income country. In early work, we have used this cohort to quantify the national HIV care cascade, to assess geographic heterogeneity in viral suppression, to assess rates of transfer across facilities, to quantify trends in clinical presentation, and to assess the shifting burden of adolescents on HIV treatment. In future work, we plan to assess the feasibility of realtime assignment of this unique identifier and utilization of the record linkage algorithm to improve patient care.

## Appendix

**Appendix Figure 1.**
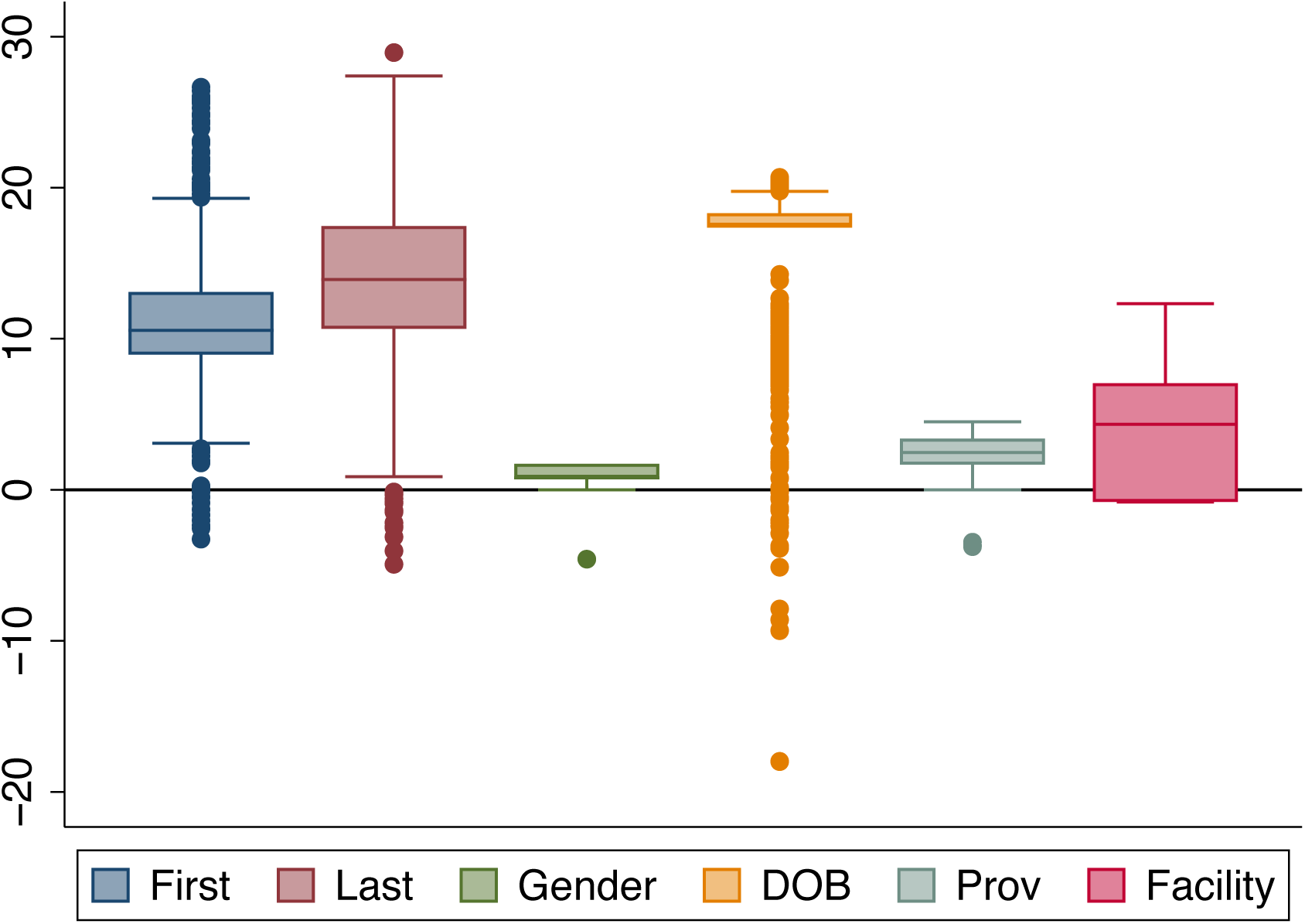
Distribution of Similarity Scores for Each Domain Among “True Matches” Identified in the Manually-Matched Training Data. Box plot shows distributions of sub-scores from each of the domains in the comparison vector: first name, last name, gender, date of birth, province, and facility. Data are limited to 3899 comparisons coded as “true matches” in the (revised) manually-matched training data. The boxes denote the interquartile range, the midline is the median, and the whiskers denote 5^th^ and 95^th^ percentiles of the distribution.

**Appendix Table 1.**
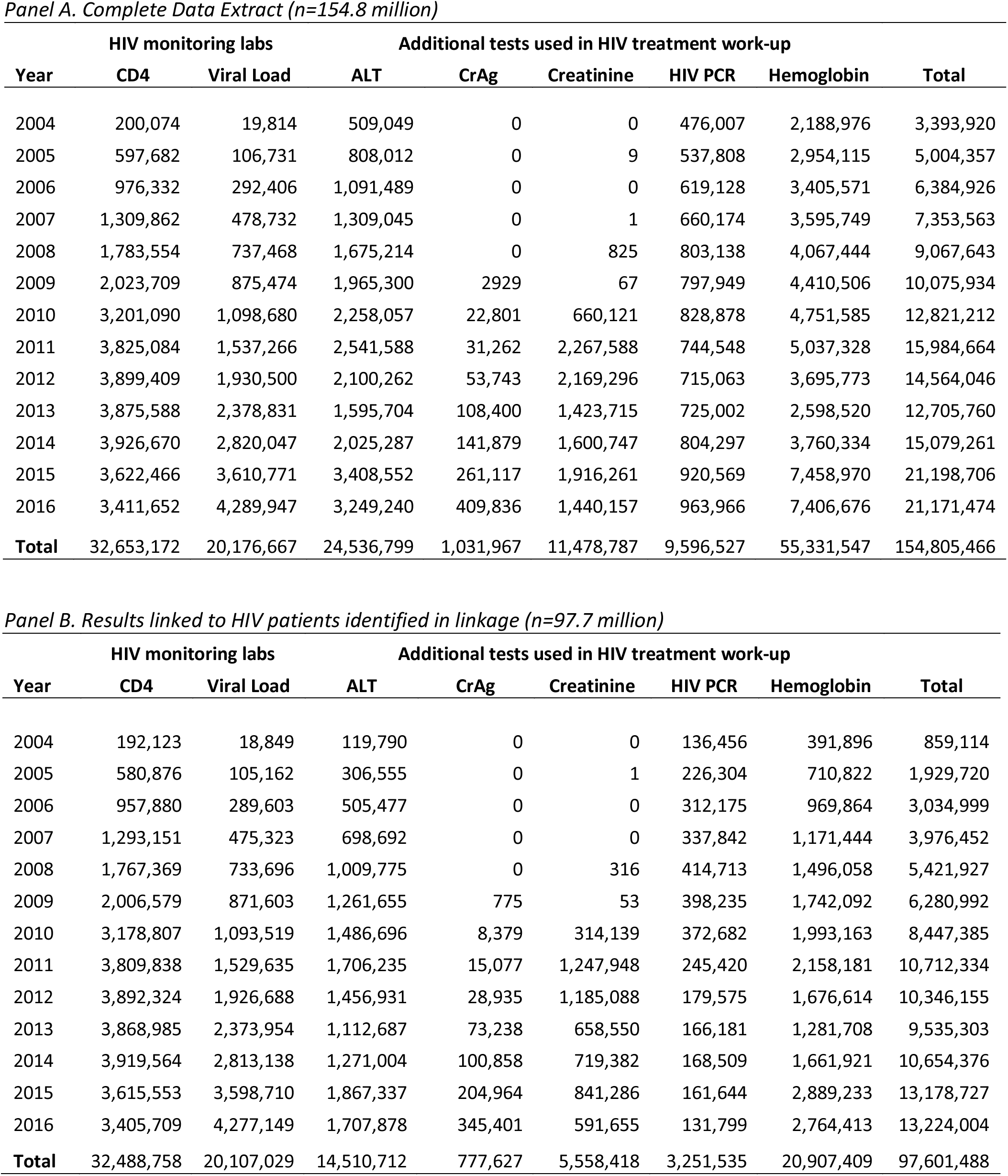
**Number of laboratory test results by type and year**

**Appendix Table 2.** **Sensitivity and PPV with 95% CIs relative to test and training data**

| Unique identifier | Sensitivity |  | Positive Predictive Value |  | F-measure |
| --- | --- | --- | --- | --- | --- |
|  | *Original* | *Corrected* | *Original* | *Corrected* | *Corrected* |
| **Exact match ID:** Test | 59.5% | 53.4% | 100.0% | 100.0% | 69.6% |
| Test 95%CI | (55-64) | (50-57) | (100-100) | (100-100) |  |
| Training | 58.8% | 55.5% | 100.0% | 100.0% | 71.4% |
| Training 95%CI | (55-63) | (52-59) | (100-100) | (100-100) |  |
| **CDW Unique ID:** Test | 71.0% | 64.0% | 99.3% | 100.0% | 78.1% |
| Test 95%CI | (67-75) | (60-68) | (99-100) | (100-100) |  |
| Training | 69.0% | 65.7% | 93.0% | 93.9% | 77.3% |
| Training 95%CI | (65-73) | (62-70) | (87-99) | (88-99) |  |
| **BU Unique ID (50):** Test | 96.5% | 96.5% | 91.5% | 97.7% | 97.1% |
| Test 95%CI | (95-98) | (95-98) | (89-93) | (96-99) |  |
| Training | 95.6% | 97.2% | 87.6% | 94.4% | 95.8% |
| Training 95%CI | (94-97) | (96-98) | (85-90) | (93-96) |  |
| **BU Unique ID (70):** Test | 95.2% | 93.7% | 93.3% | 98.6% | 96.1% |
| Test 95%CI | (93-97) | (92-96) | (91-95) | (98-100) |  |
| Training | 93.7% | 94.7% | 89.7% | 96.1% | 95.4% |
| Training 95%CI | (92-96) | (93-96) | (87-92) | (95-98) |  |
| **BU Unique ID (90):** Test | 91.9% | 88.4% | 94.8% | 99.5% | 93.6% |
| Test 95%CI | (90-94) | (86-91) | (93-97) | (99-100) |  |
| Training | 90.4% | 90.1% | 92.8% | 98.0% | 93.9% |
| Training 95%CI | (88-93) | (88-92) | (90-95) | (97-99) |  |

Note: 95% CIs are constructed via bootstrapping clusters of records associated with the reference episode\_no's randomly sampled in the manual matching exercise.

